# Inhibition of the Eukaryotic Initiation Factor-2-α Kinase PERK Decreases Risk of Autoimmune Diabetes in Mice

**DOI:** 10.1101/2023.10.06.561126

**Authors:** Charanya Muralidharan, Fei Huang, Jacob R. Enriquez, Jiayi E. Wang, Jennifer B. Nelson, Titli Nargis, Sarah C. May, Advaita Chakraborty, Kayla T. Figatner, Svetlana Navitskaya, Cara M. Anderson, Veronica Calvo, David Surguladze, Mark J. Mulvihill, Xiaoyan Yi, Soumyadeep Sarkar, Scott A. Oakes, Bobbie-Jo M. Webb-Robertson, Emily K. Sims, Kirk A Staschke, Decio L. Eizirik, Ernesto S. Nakayasu, Michael E. Stokes, Sarah A. Tersey, Raghavendra G. Mirmira

## Abstract

Preventing the onset of autoimmune type 1 diabetes (T1D) is feasible through pharmacological interventions that target molecular stress-responsive mechanisms. Cellular stresses, such as nutrient deficiency, viral infection, or unfolded proteins, trigger the integrated stress response (ISR), which curtails protein synthesis by phosphorylating eIF2α. In T1D, maladaptive unfolded protein response (UPR) in insulin-producing β cells renders these cells susceptible to autoimmunity. We show that inhibition of the eIF2α kinase PERK, a common component of the UPR and ISR, reverses the mRNA translation block in stressed human islets and delays the onset of diabetes, reduces islet inflammation, and preserves β cell mass in T1D-susceptible mice. Single-cell RNA sequencing of islets from PERK-inhibited mice shows reductions in the UPR and PERK signaling pathways and alterations in antigen processing and presentation pathways in β cells. Spatial proteomics of islets from these mice shows an increase in the immune checkpoint protein PD-L1 in β cells. Golgi membrane protein 1, whose levels increase following PERK inhibition in human islets and EndoC-βH1 human β cells, interacts with and stabilizes PD-L1. Collectively, our studies show that PERK activity enhances β cell immunogenicity, and inhibition of PERK may offer a strategy to prevent or delay the development of T1D.

## INTRODUCTION

Type 1 diabetes (T1D) is a disorder of glucose homeostasis that results from the autoimmune destruction of insulin-producing islet β cells. The importance of the immune system in initiating the early phases of T1D is emphasized in recent clinical studies showing that blockade of the T cell receptor reduces β cell stress and delays the development of T1D (1–3). These and related preclinical studies have collectively served as an impetus to shift therapeutic emphasis toward disease modification and prevention (4, 5). Molecular alterations in β cells themselves, genetically predetermined and/or initiated by environmental insults, may contribute to early disease pathogenesis by transmitting signals that initiate and/or amplify the autoimmune assault (6, 7). Environmental insults that can trigger β cell dysfunction and T1D in individuals with genetic predispositions include, among others, viral infections, systemic inflammation, and dietary factors that all alter immune tolerance (8). As a response to these insults, various stress response mechanisms such as the unfolded protein response (UPR), integrated stress response (ISR), autophagy, anti-oxidant response, and proteasomal degradation, are employed by β cells in an attempt to maintain cellular homeostasis (9).

The ISR is an evolutionarily conserved adaptive response used to mitigate cellular stress by reducing protein production burden, enhancing the expression of stress response genes such as chaperones, and inducing the degradation of misfolded proteins (10). As part of the ISR, four kinases act as sensors of distinct stress signals: PKR (induced by viral infections), PERK (induced by UPR), GCN2 (induced by nutrient deprivation), and HRI (induced by Heme deprivation). When activated, each kinase phosphorylates eukaryotic translation initiation factor 2α (eIF2α) (11), which results in sequestration of initiation factor complex eIF2B. This sequestration suppresses the translation initiation of capped mRNAs while facilitating the alternative translation of “privileged” mRNAs that serve to combat stress and promote cell survival (12, 13). The translationally-repressed mRNAs and their associated proteins aggregate to form non-membranous bodies known as stress granules, where they reside until either disassembly (post stress) or removal by autophagy (persistent stress) (14). The adaptive nature of the ISR during embryogenesis is exemplified by Wolcott-Rallison syndrome, a human disorder in which homozygous loss of function mutations in the gene encoding PERK (*EIF2AK3*) results in neonatal diabetes (15); this phenotype is mirrored in *Eif2ak3-/-* mice (16). However, the ISR may also become maladaptive, particularly in the context of disease, and thereby exacerbate disease pathogenesis. For example, heterozygous deletion of *Eif2ak3* in *Akita* mutant mice (which develop β cell loss and diabetes owing to a mutation in proinsulin that cripples its folding) significantly delays diabetes onset, a phenotype replicated by use of low-dose PERK inhibitors in these mice (17).

Whereas a maladaptive role for the β cell ISR during autoimmune T1D pathogenesis remains speculative, recent studies have shown the dysregulation of ISR genes in pancreatic tissue sections from donors with T1D and pre-T1D (18). The ISR kinase PERK is activated as one of three branches of the UPR. The roles of the other two UPR branches (ATF6 and IRE1α) have been studied in the context of T1D (19–21). Given the developmentally essential role of PERK in both the exocrine pancreas and β cell (16, 22, 23), it remains unknown if and how PERK activity might contribute to the pathogenesis of T1D. We hypothesized that prolonged activation of PERK contributes to β cell dysfunction and maintenance of autoimmunity in T1D. In this study, we utilized a novel small molecule kinase inhibitor of PERK to define the molecular effects of the ISR and its role in mouse and human T1D pathogenesis. Our findings provide evidence that the ISR, via PERK, governs a molecular response that increases susceptibility of β cells to autoimmune attack and provides a new approach to intervening during the early stages of T1D to promote disease prevention and modification.

## RESULTS

### The integrated stress response is activated in β cells of pre-diabetic NOD mice and humans

ER stress in β cells has been implicated in promoting T1D pathogenesis (24). In response to ER stress, the β cell activates the UPR, in part to reduce protein load and recover ER homeostasis. To assess UPR activation in islets in the pre-T1D period, we first reanalyzed a publicly available single-cell RNA sequencing dataset (25) of pancreatic islets isolated from female NOD mice during the pre-diabetic period (4, 8, and 15 weeks of age) (**Figure 1A**).

**Figure 1:**
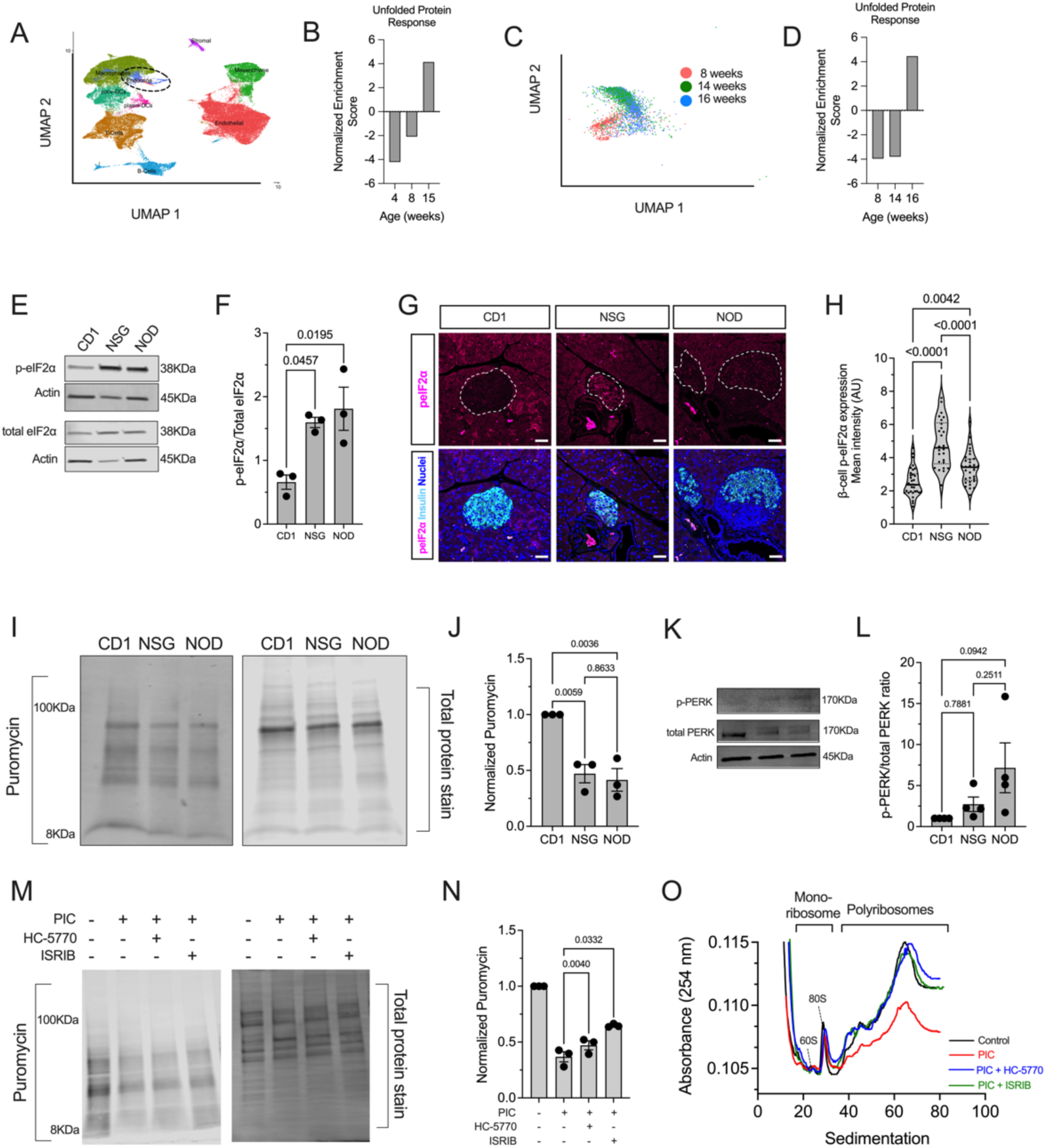
The unfolded protein response and integrated stress response are active in prediabetic NOD mice and PIC-treated human islets. (***A***) Uniform manifold approximation and projection (UMAP) embeddings of a reanalysis of single-cell RNA seq of islets from 4-, 8-, and 15-week-old female NOD mice. (***B***) Gene set enrichment analysis (GSEA) of the endocrine cell population identified in *panel* (*A*) for HALLMARK: unfolded protein response. *(**C**)* UMAP embeddings of a reanalysis of single-cell RNA seq of islets from 8-, 14-, and 16-week-old female NOD mice. (***D***) Gene set enrichment analysis (GSEA) of the β cell population identified in *panel (C)* for HALLMARK: unfolded protein response. (***E***) Representative immunoblot of phosphorylated eIF2α and total eIF2α; N=3 biological replicates. (***F***) Quantification of the immunoblots shown in *panel (E)*. ANOVA. (***G***) Representative pancreatic immunofluorescence images of phosphorylated eIF2α (magenta), insulin (cyan), and nuclei (blue); Scale bar = 50μm; *dotted lines* indicate islets. (***H***) Quantification of the fluorescence intensity of the images in *panel (G)*; each dot represents an islet, N=4-5 mice, and N>5 islets per mouse (ANOVA). (***I***) Representative puromycin incorporation immunoblot image (*left panel*) and corresponding total protein stain (*right panel*) from islets of 8-week-old female CD1, NSG, and NOD islets. (***J***) Quantification of the puromycin intensity normalized to total protein stain from *panel (I);* N=3 (each replicate represents pooled islets from 3-4 mice per group) (ANOVA). (***K***) Representative immunoblot of phosphorylated PERK and total PERK from islets of 8-week-old female CD1, NSG, and NOD mice (*top*); quantification of the immunoblot for phospho-PERK normalized to total PERK from N=4 (each replicate represented pooled islets from 3-4 mice per group) (*bottom*) (ANOVA) (L) Representative puromycin incorporation immunoblot (*left panel*) and corresponding total protein stain (*right panel*) from MIN6 cells treated as indicated (PIC=proinflammatory cytokines). (*M*) Quantification of the puromycin intensity normalized to total protein stain from panel (*L*); N=3 independent experiments (RM-ANOVA). (***N***) Polyribosomal profiling traces of human islets treated ±PIC, HC-5770, or ISRIB. Data are presented as mean ±SEM.

Because the dataset was enriched for immune cell populations, our analysis focused on the endocrine cell subset (composed largely of β cells) without stratifying individual cell types. Gene set enrichment analysis (GSEA) revealed a gradual enrichment of genes of the UPR with advancing age in the endocrine cell population (**Figure 1B**). To specifically assess β cells, we reanalyzed a different publicly available single-cell RNA sequencing dataset (26) of islets from pre-diabetic female NOD mice (8, 14, and 16 weeks of age) (**Figure 1C**). Similar to our observation with the endocrine cell population, GSEA showed an enrichment of the UPR over time in β cells (**Figure 1D**). Because the UPR includes a molecular arm (via PERK) that activates the ISR, we next probed for a hallmark feature of the ISR, phosphorylated (Ser51)- eIF2α (p-eIF2α) in pancreatic tissue of female NOD mice between 6-12 weeks of age.

Relatively higher p-eIF2α immunostaining in β cells was observed in 6- and 8-week-old mice, compared to 10- and 12-week-old mice (**Supplemental Figure 1A and B**), suggesting that the ISR is maximally activated before 10 weeks of age in NOD mice. Further studies were therefore focused on NOD mice before 10 weeks of age. Because male NOD mice develop diabetes at a lower frequency than females, we compared the relative p-eIF2α immunostaining in males and females at 8 weeks of age but observed no differences (**Supplemental Figure 1C and D**). This finding suggests that activation of the ISR at this early age does not account for differences in diabetes incidence between the sexes.

To assess the relative activity of the ISR in NOD mice compared to controls, islets and tissues from 8-week-old female NOD mice were compared to age- and sex-matched, diabetes-resistant CD1 and NSG (*NOD-scid IL-2R-γ-null*) mice. NSG mice share the genetic background of the NOD mice but lack a functional immune system and therefore do not develop diabetes.

Immunoblotting of isolated islets showed increased p-eIF2α levels in NOD and NSG islets compared to CD1 islets (**Figure 1E and F**). Because islets from immunodeficient NSG mice also show an increase in p-eIF2α levels, this finding emphasizes that activation of the ISR is a feature of islets on the NOD genetic background that is independent of the immune system. To confirm ISR activation in β cells specifically, we performed immunofluorescence staining for p- eIF2α in tissues from 8-week-old mice. Both NOD and NSG mice showed an increase in p- eIF2α immunostaining compared to CD1 controls (**Figure 1G and H**). Together, these data suggest that the ISR in β cells is activated before the onset of overt hyperglycemia and may be a feature independent of the immune system in T1D-prone NOD mice.

### The ISR induces global mRNA translational initiation blockade

The phosphorylation of eIF2α during the ISR leads to sequestration of the translation initiation factor eIF2B, suppressing mRNA translation initiation (for a review see (27)) and thereby reducing protein synthesis. To assess the effects of ISR on protein synthesis, we performed surface sensing of translation (SUnSET) (28) on isolated islets from 8-week-old CD1, NSG, and NOD mice. Incorporation of puromycin into elongating polypeptide chains, followed by immunoblotting with anti-puromycin antibodies, allows for assessment of mRNA translation. Consistent with the activation of the ISR, we observed reduced puromycin incorporation into proteins of NSG and NOD islets, suggesting that global mRNA translation is reduced in pre- diabetic stages (**Figure 1I and J**). ISR activation can be mediated by any one or more of the four kinases—PERK, PKR, GCN2, and HRI. Based on our observed increase in UPR in NOD islets over time (**Figure 1A-D)**, we surmised that PERK may be the relevant activated kinase in

NOD islets. Immunoblot analysis demonstrated an increased trend in phosphorylated PERK in islets of both NSG and NOD mice compared to CD1 controls, implicating PERK as the ISR kinase (**Figure 1K and L**). We next performed SUnSET using mouse MIN6 β cells treated with proinflammatory cytokines (IFN-γ + IL-1β + TNF-α) to induce a state mimicking β cell ER stress seen in T1D (29). Like NOD islets, we observed reduced puromycin incorporation into proteins of proinflammatory cytokine-treated cells compared to vehicle control (**Figure 1M**). To directly correlate the block in protein synthesis with the activity of the ISR, we utilized two inhibitors of the ISR. HC-5770 is a highly specific inhibitor of PERK (previously characterized as Cmpd26 in (30)), and ISRIB is a previously described inhibitor of the p-eIF2α/eIF2B interaction (31). Co-treatment with 250 nM HC-5770 or 50 nM ISRIB partially reversed the block in protein synthesis induced by proinflammatory cytokines (**Figure 1M and N**).

To interrogate the effects of the ISR on mRNA translation initiation, we performed polyribosome profiling (PRP) studies of total RNA from cadaveric human donor islets. PRP can distinguish global changes in mRNA translation initiation and elongation by assessment of the RNA sedimenting with polyribosomes vs. monoribosomes (**Figure 1O**). Lower polysome sedimentation of RNA suggests a relative translation initiation blockade (32). Human islets treated with proinflammatory cytokines (IFN-γ + IL-1β) to mimic T1D inflammation (29) showed reduced association of RNA with polyribosomes (or translation initiation blockade) by PRP compared to control islets (**Figure 1O**). Concurrent treatment of human islets with either 250 nM HC-5770 or 50 nM ISRIB led to a recovery of the RNA sedimenting with polysomes, reversing the effects of proinflammatory cytokines (**Figure 1O**). Collectively, these data indicate that inflammation induces a translation initiation blockade, which is reversed upon inhibition of the ISR.

### Pharmacokinetic and pharmacodynamic assessment of the PERK inhibitor HC-5770

To evaluate the role of PERK in β cell dysfunction in vivo, we utilized the PERK inhibitor HC-5770. First, to ensure that HC-5770 does not cause a compensatory activation of GCN2 (as reported for other PERK inhibitors) (33), we performed a dose-ranging study in isolated CD1 mouse islets followed by immunoblotting for phospho-GCN2 and total GCN2. Between 8-1000 nM concentrations of HC-5770, we observed no changes in phospho-GCN2 (**Supplemental Figure 2A**). To ensure that our antibody detects phospho-GCN2, we also performed immunoblotting of mouse embryonic fibroblasts (MEFs), which showed that 100 nM halofuginone (a GCN2-ISR activator) (34) robustly induces p-GCN2, while thapsigargin treatment suppresses it (**Supplemental Figure 2A**). Initial pharmacokinetic and pharmacodynamic (PK/PD) analyses were performed to confirm in vivo PERK inhibition in mouse pancreas and to identify appropriate doses for further study. The first PK analysis of HC- 5770 followed a single oral administration of HC-5770 at doses ranging from 0.3 to 30 mg/kg in BALB/c mice, which revealed dose-proportionate increases in plasma exposure with a half-life of approximately three hours (**Table 1**). The unbound fraction (Fu) in mouse plasma was determined in vitro to be 0.3%, which enabled us to calculate the free, unbound drug plasma exposure across time in vivo (**Supplemental Figure 2B**). A second PK experiment in NOD mice followed a single oral administration of HC-5770 at 1 and 10 mg/kg and confirmed nearly identical exposure and clearance between NOD and BALB/c mouse strains (**Supplemental Figure 2C**).

**Table 1:** PK analysis of HC-5770 in mouse plasma and pancreas. HC-5770 quantified by LC-MS/MS following single oral administration at doses ranging from 0.3 to 30 mg/kg in BALB/c mice (n=5 mice/group/timepoint).

|  | Plasma |  |  |  | Pancreas |  |  |  |
| --- | --- | --- | --- | --- | --- | --- | --- | --- |
| **Dose (mpk)** | **C<sub>max</sub> (ng/ml)** | **Terminal t<sub>1/2</sub> (h)** | **AUC<sub>last</sub> (h\*ng/mL)** | **AUC<sub>INF</sub> (h\*ng/mL)** | **C<sub>max</sub> (ng/ml)** | **Terminal t<sub>1/2</sub> (h)** | **AUC<sub>last</sub> (h\*ng/mL)** | **AUC<sub>INF</sub> (h\*ng/mL)** |
| **0.3** | 306 | 2.8 | 1754 | 1761 | 91 | 3.5 | 479 | 535 |
| **1** | 1239 | 3 | 6830 | 6865 | 355 | 3.2 | 2170 | 2185 |
| **3** | 3838 | 2.9 | 26094 | 26245 | 1314 | 3 | 8547 | 8600 |
| **10** | 11072 | 3.4 | 103631 | 104712 | 3309 | 3.5 | 35828 | 36265 |
| **30** | 27920 | 5.3 | 321718 | 339265 | 9420 | 5 | 118125 | 123646 |

The PD effect of HC-5770 on phosphorylated PERK (T980; pPERK) was evaluated in mouse pancreas. Whole protein lysates from mouse pancreata isolated from BALB/c mice following single administration of HC-5770 at doses ranging from 0.3 to 30 mg/kg, as described above. At 10 and 30 mg/kg, HC-5770 achieved approximately 75% inhibition 1 h post-dose, whereas doses ranging from 0.3 – 3 mg/kg induced moderate effects on pPERK/PERK levels that were sustained past 4 h following administration (**Supplemental Figure 2D**). We next sought to evaluate the impact of PERK inhibition on insulitis by treating prediabetic NOD mice with HC-5770 at doses ranging from 0.3 to 30 mg/kg (twice daily) for two weeks. Following the treatment period, mouse pancreas sections were stained and scored for the level of islet immune infiltration (insulitis). HC-5770 decreased insulitis at all doses tested, with the greatest response noted at doses of 1 mg/kg and above (**Supplemental Figure 2E**). As complete and sustained PERK inhibition has previously been associated with pancreatic degeneration and dysfunction (35), we reasoned that the lowest efficacious doses should be selected for continued investigation in vivo and selected 0.3, 1, and 3 mg/kg twice daily as reasonable doses to advance. Flexibility in the dosing regimen was then evaluated by assessing the insulitis response in animals treated twice daily vs. once daily with HC-5770. NOD mice were treated either twice daily at 0.3, 1, and 3 mg/kg or once daily at 0.6, 2, and 6 mg/kg HC-5770 for two weeks. The once-daily dosing schedules resulted in similar effects as twice-daily dosing on insulitis, significantly inhibiting insulitis at all doses tested (**Supplemental Figure 2F**). Based on these findings, once-daily doses ranging from 0.6 to 6 mg/kg (QD) were selected for continued evaluation in vivo.

### Systemic inhibition of PERK delays autoimmune diabetes development in NOD mice

We hypothesized that the β cell translational blockade induced by the ISR in the pre- diabetic state is maladaptive and contributes to the development of T1D. To test this hypothesis, we treated female NOD mice with HC-5770 for 4 weeks during the pre-diabetic stage when the ISR is active (6-10 weeks of age) and monitored for subsequent diabetes development until 25 weeks of age (see scheme in **Figure 2A**). NOD mice were treated with either vehicle or three different doses of HC-5770 (0.6, 2, or 6 mg/kg per day). Approximately 60%, 53%, and 68% of the mice treated with 0.6 mg/kg, 2 mg/kg, and 6 mg/kg of HC-5770, respectively, remained diabetes-free by 25 weeks of age, whereas only 10% of the vehicle-treated mice remained diabetes-free (**Figure 2B**). These findings suggest an enduring effect of early PERK inhibitor treatment. Notably, the exocrine pancreas of mice treated with HC-5770 (6 mg/kg) showed no gross pathological evidence of pancreatitis (**Figure 2C**), unlike findings observed upon more complete and sustained inhibition with other PERK inhibitors (36). This observation is also supported by gross pancreas weights that were unaffected by two weeks of treatment with either the twice-daily or once-daily dosing regimens (**Supplemental Figure 2G**).

**Figure 2:**
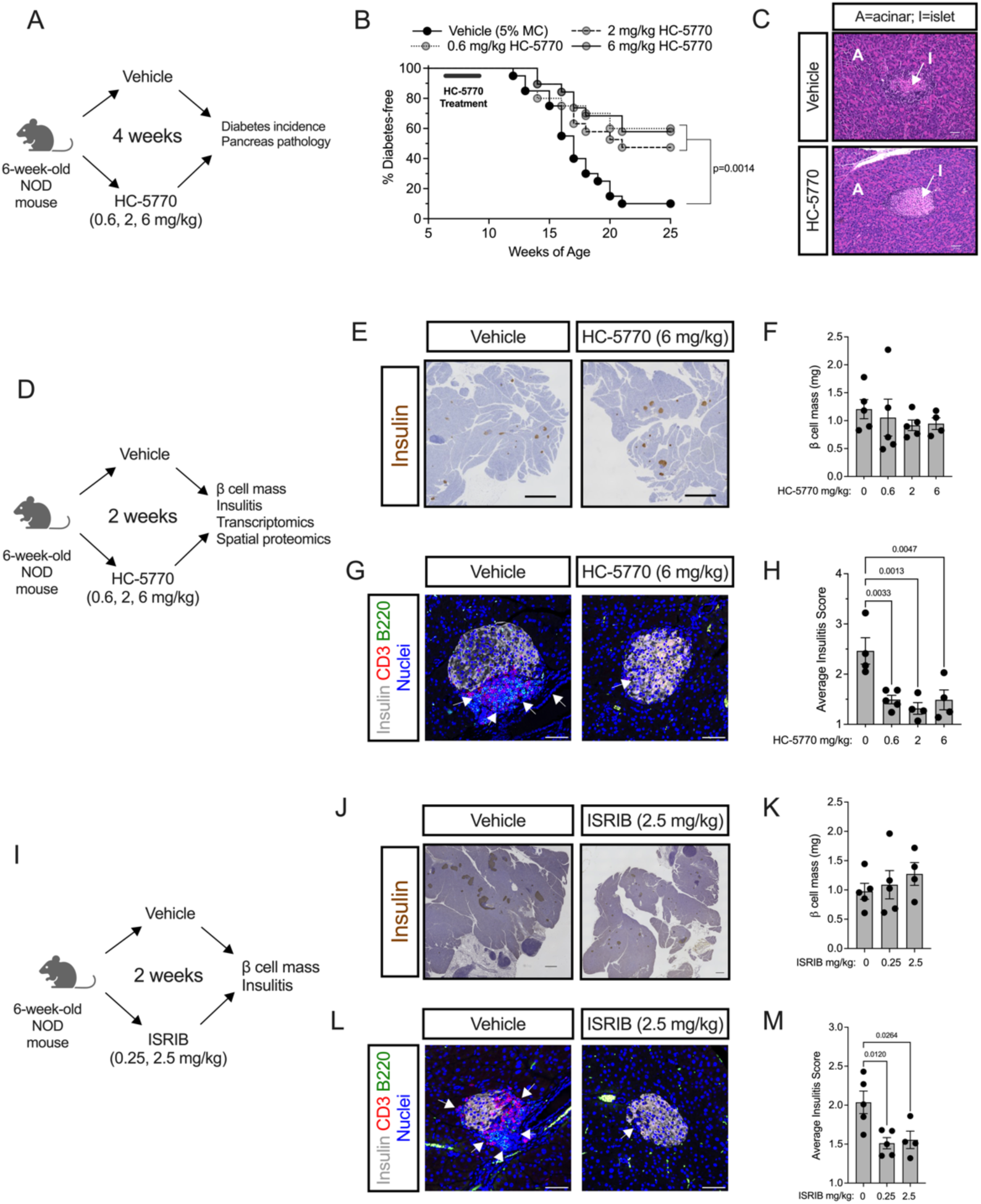
PERK inhibition delays autoimmune diabetes in NOD mice. Prediabetic female NOD mice (6 weeks of age) were treated with varying doses of HC-5770 or ISRIB. (***A***) Experimental design for HC-5770 diabetes incidence study. (***B***) Diabetes incidence. N=20 mice per group (Mantel-Cox). (***C***) Representative hematoxylin and eosin stain of pancreata from non-diabetic mice at 25 weeks of age that were treated with Vehicle or 6 mg/kg HC-5770 from 6-10 weeks of age. (***D***) Experimental design for HC-5770 mechanistic short-term studies. (***E***) Representative images of pancreata from NOD mice following two weeks of HC-5770 administration (6 mg/kg) stained for insulin (*brown*) and counterstained with hematoxylin (*blue*); scale bar = 500 μm. (***F***) β cell mass of mice treated with HC-5770 (6 mg/kg) for two weeks; N=4-5 mice per group (ANOVA). (***G***) Representative images of pancreata from NOD mice following two weeks of HC-5770 administration (6 mg/kg) immunostained for CD3 (*red*), B220 (*green*), insulin (*white*), and nuclei (*blue*); *arrows* indicate regions of insulitis; scale bar = 50 μm. (***H***) Average insulitis score of mice treated with varying doses of HC-5770 for two weeks; N=4-5 mice per group; NB: these data are replicated in Supplemental Figure 2F for comparative purposes (ANOVA). (***I***) Experimental design for ISRIB short-term studies. (***J***) Representative images of pancreata from NOD mice following two weeks of ISRIB administration (2.5 mg/kg) stained for insulin (*brown*) and counterstained with hematoxylin (*blue*); scale bar = 500 μm. (***K***) β cell mass of mice treated with ISRIB (2.5 mg/kg) for two weeks; N=4-5 mice per group (ANOVA). (***L***) Representative images of pancreata from NOD mice following two weeks of ISRIB administration (2.5 mg/kg) immunostained for CD3 (*red*), B220 (*green*), insulin (*white*), and nuclei (*blue*); *arrows* indicate regions of insulitis; scale bar = 50 μm. (***H***) Average insulitis score of mice treated with varying doses of ISRIB for two weeks; N=4-5 mice per group (ANOVA). Data are presented as mean ±SEM.

### HC-5770 treatment engages molecular pathways related to PERK functions in β cells, reduces β cell death, and enhances β cell replication

To identify proximal molecular effects of PERK inhibition and its impact on the islet microenvironment, we next performed a short-term, 2-week oral treatment of pre-diabetic NOD mice (beginning at 6 weeks of age) with differing doses of HC-5770 followed by assessment of glucose homeostasis, pancreas pathology, and islet single cell molecular analyses (see scheme in **Figure 2D**). Upon treatment at any of the HC-5770 doses, there was no statistical change in β cell mass (**Figure 2E and F**) compared to controls. However, as noted previously, there was a significant decrease in insulitis in HC-5770-treated mice at all doses compared to vehicle controls (**Figure 2G and H**), a finding preceding the eventual protection of these mice from diabetes. To confirm that the PERK inhibition effect occurs via blockade of p-eIF2α function, we next utilized ISRIB (an inhibitor of the p-eIF2α/eIF2B interaction) in NOD mice. Six-week-old NOD mice were treated with two different doses of ISRIB (0.25 or 2.5 mg/kg) or vehicle by intraperitoneal injection for 2 weeks (see scheme in **Figure 2I**). Consistent with the effects of HC-5770, mice receiving ISRIB exhibited no effect on β cell mass (**Figure 2J and K**) and a significant reduction in insulitis (**Figure 2L and M**).

To determine the effect of HC-5770 on molecular pathways in the cells of the islet microenvironment, we performed single-cell RNA sequencing (scRNA-seq) of islets following 2- week oral treatment of NOD mice beginning at 6 weeks of age. For these studies, we employed HC-5770 at 6 mg/kg, as the mice that received this dose in our diabetes outcome study had the lowest incidence of diabetes. We visualized cells based on expression profiles using uniform manifold approximation and projection (UMAP) for dimension reduction plots and identified clusters representing distinct pancreatic cell types (**Figure 3A**). β, α, δ, PP, acinar, stellate, duct, T cells, B cells, and myeloid cell types were characterized based on expression of genes *Ins1/2, Gcg, Sst, Ppy, Prss1, Col3a1, Krt19, Trbc2, Cd79a, H2-Eb1*, respectively. Dot plots of the top 5 genes in each cell type confirm the correct identification of cell types (**Supplemental Figure 3A**).

**Figure 3:**
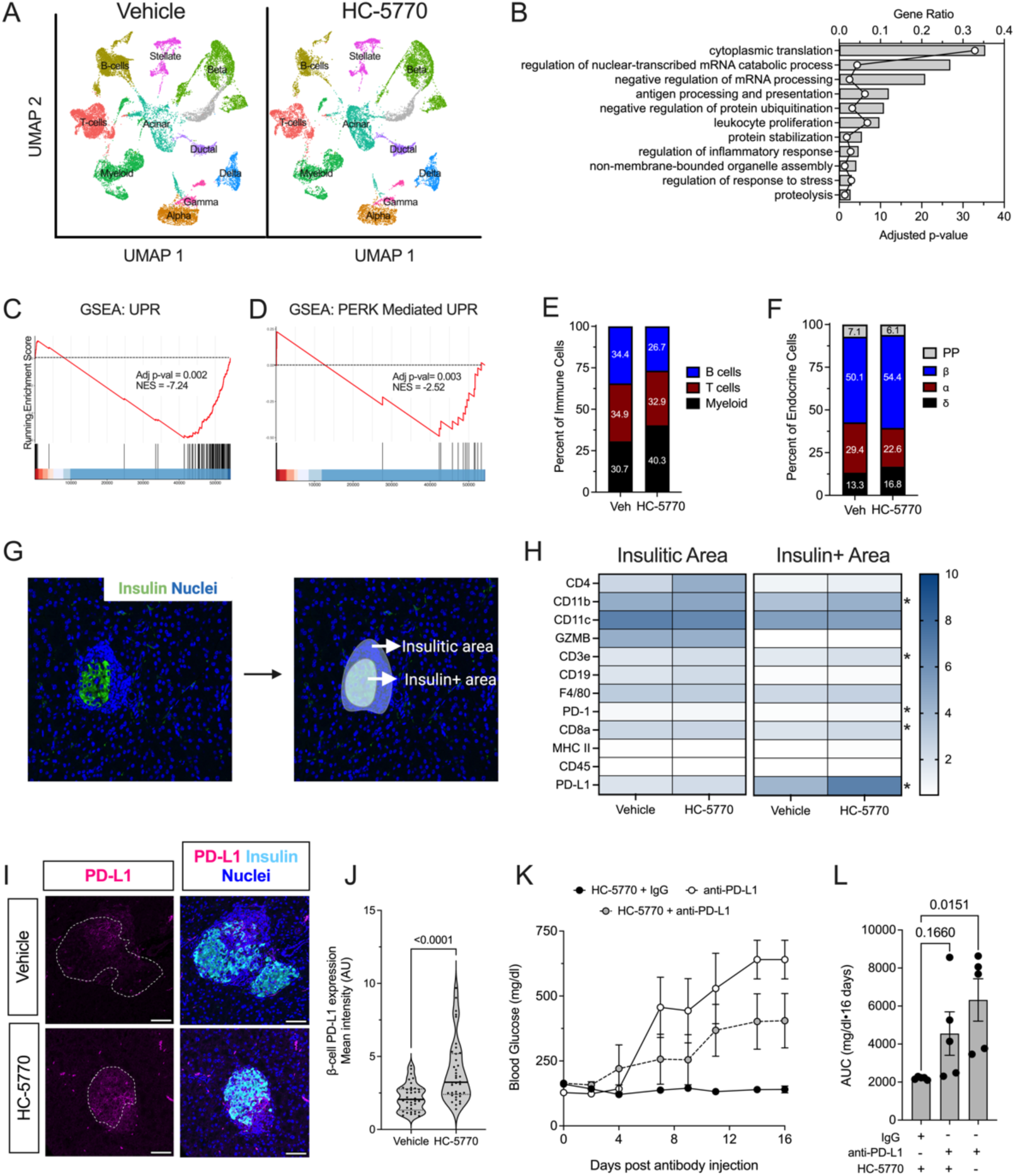
PERK inhibition increases levels of PD-L1 in β cells of NOD mice. Prediabetic female NOD mice (6 weeks of age) were treated with 6 mg/kg HC-5770 for 2 weeks and isolated islets were subjected to scRNA-seq and pancreas tissue was subjected to Nanostring® spatial proteomics. (***A***) Uniform manifold approximation and projection (UMAP) embeddings of merged single-cell RNA sequencing profiles from islets colored by identified cell clusters. N=3 mice per group for scRNA-seq. (***B***) Gene Ontology analysis of all cell clusters (pseudo-bulk analysis). (***C***) Gene set enrichment analysis (GSEA) of β cell clusters showing HALLMARK: unfolded protein response and (***D***) GSEA of β cell clusters showing GO-BP: PERK-meditated unfolded protein response. (***E***) Percent of T, B, and myeloid cells identified within the immune cell clusters. (***F***) Percent of α, β, δ, and PP cells identified within the islet cell clusters. (***G***) Representative image of identification of the insulin-positive area and the insulitic area used to collect spatial tissue-based proteomics. (***H***) Heatmap of identified proteins in the insulitic area (*left panel*) and insulin-positive area (*right panel*) in Vehicle- or HC-5770-treated mice; N=10-11 regions of interest from 2 mice per group. * indicates P<0.05 (T-test). (***I***) Representative images of pancreata from mice following two weeks of treatment with Vehicle or HC-5770 stained for PD-L1 (*magenta*), insulin (*cyan*), and nuclei (*blue*); *dotted lines* indicate islets; scale bar=50 μm. (***J***) Quantification of PD-L1 intensity in the β cells of the panel (I); each dot represents an islet, N=4-5 mice, and N>5 islets per mouse (T-test). (***K***) 6-week-old female NOD mice were treated as indicated with 6 mg/kg HC-5770 or with Vehicle for two weeks, then administered either anti-PD-L1 or IgG control, followed by another two weeks of HC-5770 treatment until 10 weeks of age, and glucoses were measured on alternate days post-injection; N= 5 mice per group. (***L***) Area Under the Curve (AUC) analysis of the data in *panel (K)* (ANOVA). Data are presented as mean ±SEM.

To assess the engagement of molecular processes by HC-5770, we performed a pseudo-bulk analysis followed by Gene Ontology (GO) analysis. We found that cytoplasmic translation, leukocyte proliferation, digestion, protein stabilization, and protein ubiquitination were among the top significantly regulated pathways in HC-5770-treated islets (**Figure 3B**). Furthermore, we performed GO analysis on β cell clusters (9 clusters). We found that cytoplasmic translation, protein folding, ER stress response, and antigen processing and presentation were among the top significantly regulated pathways in HC-5770-treated β cells (**Supplemental Table 1**). Consistent with these findings, GSEA showed that β cells of HC- 5770-treated mice downregulated genes in the UPR pathway relative to vehicle controls, as indicated by a normalized enrichment score (NES) of -7.24 (**Figure 3C**). Because the UPR pathway encompasses three distinct arms—PERK, IRE1α and ATF6—we analyzed the PERK pathway by GSEA (prioritizing the genes *Atf4, Eif2s1, Eif2ak3, Nck1, Nck2, Nfe2l2, Ptpn1, Ptpn2, Agr2, Abca7, Bok, Tmed2, Tmem33, Qrich1*). This analysis revealed a significant decrease in PERK-mediated UPR in the β cells of HC-5770-treated mice compared to vehicle controls (NES: -2.52) (**Figure 3D**). Collectively, these data support the suppression of PERK-related molecular processes by HC-5770, indicating appropriate target engagement.

Examination of cell clusters and numbers in the HC-5770 group compared to vehicle controls revealed several notable findings: (a) there was a decrease in the percentage of T and B cells, with an increase in the proportion of myeloid cells (**Figure 3E**); (b) GO analysis (**Supplemental Figure 3B-D**) of T, B, and myeloid cells revealed alterations in cytoplasmic translation and T cell proliferation (T cells), cytoplasmic translation and the humoral immune response (B cells), and antigen processing and chemotaxis (myeloid cells)—all indicating phenotypic changes in these immune cell populations that likely influenced their function in response to PERK inhibition; and (c) with respect to T cells, in particular, GSEA analysis revealed a reduction in T cell activation (normalized enrichment score of -11.8) upon PERK inhibition (**Supplemental Figure 3E**). Accordingly, the β cell population demonstrated a reduced inflammatory response by GSEA (normalized enrichment score -6.85) (**Supplemental Figure 3F**).

Regarding the endocrine cell population, there was an increase in the overall percentage of β cells and a decrease in α cell percentage (**Figure 3F**). The increased β cell numbers upon HC-5770 treatment led us to investigate β cell replication and death. Immunostaining of pancreata showed a trend towards an increased number of proliferating cell nuclear antigen (PCNA)-positive β cells upon HC-5770 treatment (**Supplemental Figure 3G and H)**, and no changes in β cell death by terminal deoxynucleotidyl transferase dUTP nick end labeling (TUNEL) assay (**Supplemental Figure 3G and I**). In addition, scRNA-seq revealed an increase in the percentage of β cells in S/G2M phases consistent with the increased trend in PCNA-positive β cells (**Supplemental Figure 3J**).

### PERK inhibition increases β cell PD-L1 levels

To interrogate the nature of the immune cell populations in the islet microenvironment in PERK inhibitor-treated NOD mice, we performed spatial tissue-based proteomics after 2 weeks of HC-5770 treatment. We used insulin immunostaining and nuclei staining to identify β cells and the surrounding insulitic regions, respectively (**Figure 3G**). Pre-validated antibodies in the GeoMx^®^ mouse immune panel were used to probe for immune cell subtypes in the peri-islet insulitic region and within the islet. Whereas there were no statistical differences in the immune cell subtype populations in the insulitic regions of HC-5770-treated mice versus vehicle controls (**Figure 3H**), within the β cell region, there was a striking and significant upregulation of programmed death-ligand 1 (PD-L1) as well as elevations of its cognate receptor PD-1, CD3e, CD8a, and CD11b (**Figure 3H**) following PERK inhibition. The increase in PD-L1 levels on β cells was confirmed by immunofluorescence staining of pancreatic tissues (**Figure 3I and J**). The interaction of PD-L1 on β cells with PD-1 on immune cells is known to skew immune cell populations to a more immunosuppressive phenotype (37). To assess if increased PD-L1 levels after PERK inhibition mediate protection against T1D development, we treated female NOD mice that received HC-5770 for two weeks (6-8 weeks of age) with a single dose of monoclonal antibody against PD-L1 or the corresponding isotype IgG control, then continued HC-5770 treatment for an additional two weeks. As a control for diabetes development, 8-week-old female NOD mice were treated with a single dose of anti-PD-L1. HC-5770-treated mice that received isotype IgG control remained normoglycemic, while mice that received only anti-PD-L1 became hyperglycemic within 7 days (**Figure 3K and L**). The mice receiving both HC-5770 and anti-PD-L1 displayed average glucose levels intermediate between those of the two controls (**Figure 3K and L**), emphasizing that anti-PD-L1 administration partially antagonized the effect of HC-5770.

### Augmentation of PD-L1 levels requires post-translational stabilization by Golgi membrane protein 1

We next sought to clarify the molecular link between the ISR and PD-L1 levels in β cells. We first interrogated a proteomics dataset previously published by our group, in which EndoC- βH1 human β cells were treated with proinflammatory cytokines (IL-1β + IFN-γ) or vehicle (38). Proteins significantly increased following cytokine treatment included PD-L1 and Golgi membrane protein 1 (GOLM1) (**Figure 4A**). GOLM1 is a Golgi-associated protein that functions, in part, as a chaperone for protein trafficking (39) and has been shown in hepatocellular carcinoma to positively regulate PD-L1 production (40). The increase in GOLM1 protein levels following cytokine treatment was confirmed by immunoblotting in EndoC-βH1 cells (**Figure 4B**) and seen as a trend in primary human islets (**Supplemental Figure 4A**). In EndoC-βH1 cells, the increase in PD-L1 protein levels following cytokine treatment appears to be a transcriptional response, as both *GOLM1* and *CD274* (encoding PD-L1) mRNA levels increased following cytokine treatment (**Figure 4C and D).** However, in human islets, only the increase in PD-L1 protein levels following cytokine treatment appeared to be a transcriptional response (**Supplemental Figure 4B**), as no significant increase in *GOLM1* mRNA levels was observed (**Supplemental Figure 4C**). Notably, the additional increase in PD-L1 protein levels seen with PERK or ISR inhibition was not associated with substantial increase in *CD274* or *GOLM1* mRNA in EndoC-βH1 cells (**Figure 4C and D**) or human islets (**Supplemental Figure 4B and C**), suggesting that upregulation of PD-L1 with PERK/ISR inhibition is a post-transcriptional process.

**Figure 4:**
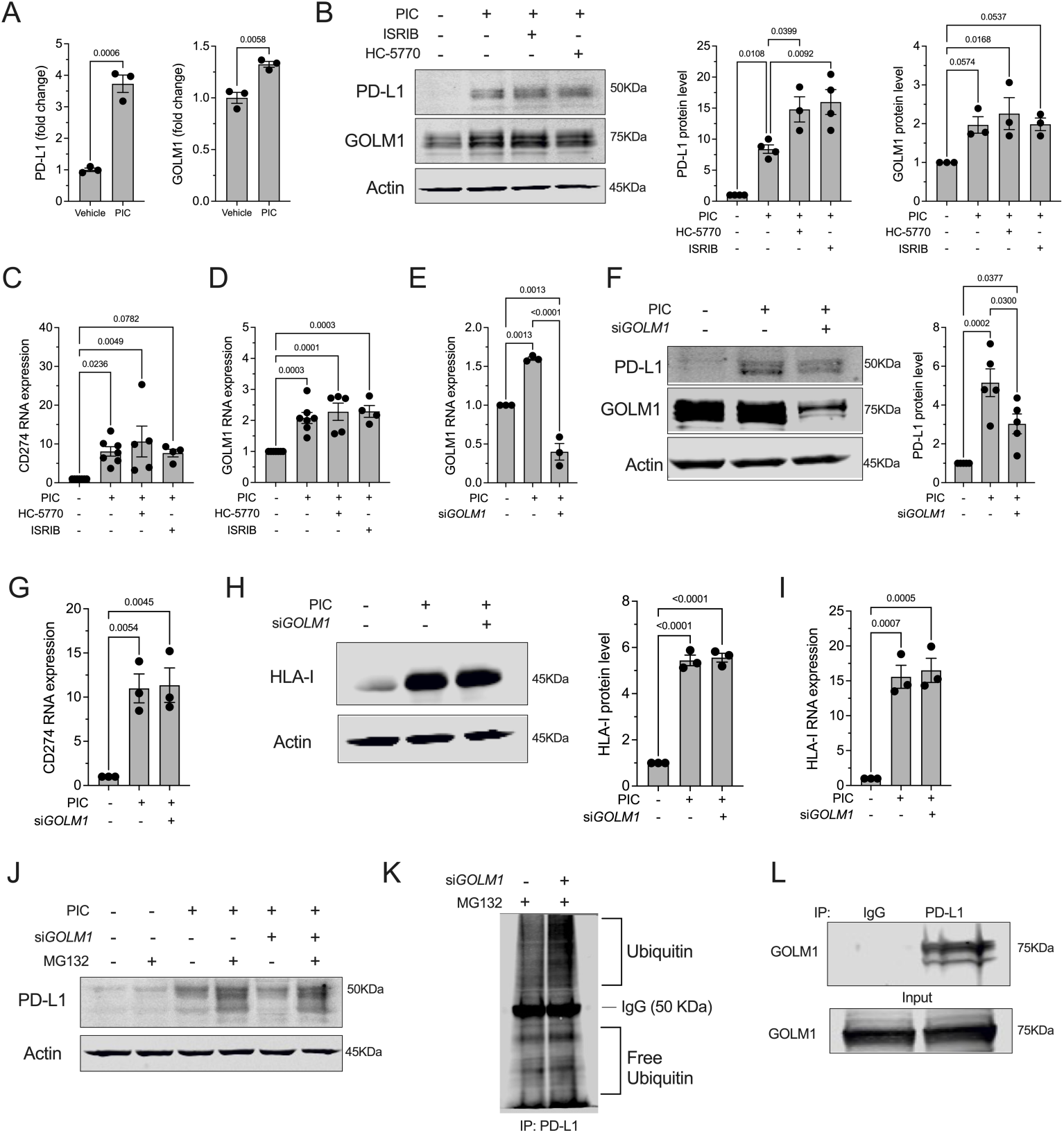
GOLM1 stabilizes PD-L1. (***A***) PD-L1 and GOLM1 protein levels were identified using proteomics of EndoC-βH1 human β cells treated ±proinflammatory cytokines (PIC); N=3 biological replicates. T-test. (***B***) Representative immunoblot analysis of PD-L1 and GOLM1 from EndoC-βH1 cells treated ±PIC, HC-5770, and ISRIB (*left panel*) with quantification of PD-L1 levels (*middle panel*) and GOLM1 levels (*right panel*); N=3 biological replicates (ANOVA). (***C, D***) Relative *CD274* and *GOLM1* mRNA levels by quantitative RT-PCR normalized to *ACTB* from EndoC-βH1 cells treated ±PIC, HC-5770, and ISRIB; N=4-7 biological replicates (ANOVA). (***E***) Relative *GOLM1* RNA levels normalized to *ACTB* from EndoC-βH1 cells treated ±PIC and *GOLM1* siRNA; N=3 biological replicates (ANOVA). (***F***) Representative immunoblot analysis of PD-L1 and GOLM1 from EndoC-βH1 cells treated ±PIC and *GOLM1* siRNA (*left panel*) with quantification of PD-L1 levels (*right panel*); N=3 biological replicates (ANOVA). (***G***) Relative *CD274* mRNA levels by quantitative RT-PCR normalized to *ACTB* of EndoC-βH1 cells treated ±PIC and *GOLM1* siRNA; N=3 biological replicates (ANOVA). (***H***) Representative immunoblot analysis of HLA-I from EndoC-βH1 cells treated ±PIC and *GOLM1* siRNA (*left panel*) with quantification of HLA-I levels (*right panel*). N=3 biological replicates (ANOVA). (***I***) Relative *HLA-I* mRNA levels by quantitative RT-PCR normalized to *ACTB* from EndoC-βH1 cells treated ±PIC and *GOLM1* siRNA; N=3 biological replicates (ANOVA). (***J***) Representative immunoblot analysis of PD-L1 from EndoC-βH1 cells treated ±PIC, *GOLM1* siRNA, and MG132. (***K***) Immunoblot analysis of ubiquitin following immunoprecipitation (IP) for PD-L1 from HEK-293 cells treated with *GOLM1* siRNA ±MG132. (***L***) Immunoblot analysis for GOLM1 following IP for PD-L1 from HEK-293 cells. Data are presented as mean ±SEM.

To investigate the potential dependence of PD-L1 production on GOLM1, we next performed siRNA-mediated silencing of *GOLM1* in EndoC-βH1 cells (**Figure 4E**). Upon GOLM1 silencing, PD-L1 protein levels, but not its encoding *CD274* gene levels, were significantly attenuated with cytokine treatment (**Figure 4F and G**), suggesting that GOLM1 is required for the maintenance of PD-L1 protein levels. By contrast, GOLM1 is not required for the production or maintenance of another known cytokine-induced molecule in β cells, human leukocyte antigen I (HLA-I) (**Figure 4H and I**)—signifying that GOLM1 does not function to promote production of all cytokine-responsive proteins.

PD-L1 protein levels are known to be regulated by post-translational modification (glycosylation and ubiquitination) (41). We tested the possibility that GOLM1 might affect PD-L1 protein stability by preventing its turnover by the proteasome. The attenuation of PD-L1 levels upon GOLM1 knockdown was partially reversed upon concurrent treatment of cells with MG132, an inhibitor of proteasome-mediated degradation (**Figure 4J**). This finding suggests that GOLM1 stabilizes PD-L1, preventing its sequestration by the proteasome. Consistent with this finding, we observed (a) that GOLM1 knockdown increases PD-L1 ubiquitination in HEK-293 cells (**Figure 4K**) and (b) there is a physical interaction between PD-L1 and GOLM1 based on co-immunoprecipitation studies in transfected HEK-293 cells (**Figure 4L**). These observations are in agreement with the decrease in ubiquitin-proteasome degradation pathway in the β cells of islets treated with HC-5770 from our scRNA-seq studies (**Supplemental Figure 3D**).

To identify the post-transcriptional mechanism whereby the ISR regulates GOLM1 and PD-L1 levels, we quantified the mRNA levels of *CD274* (encoding PD-L1) and *GOLM1* in the polyribosome and monoribosome fractions of human islets that were treated with cytokines in the presence or absence of HC-5770 or ISRIB (from **Figure 1N**). We observed no significant change in *CD274* mRNA or *GOLM1* mRNA in the polyribosome fraction (actively translating) relative to the monoribosome fraction following proinflammatory cytokine treatment (**Supplemental Figure 4E and F**). In the case of *CD274*, this finding suggests that the increased PD-L1 levels following cytokine treatment is likely due to an increase in *CD274* transcript levels. In the case of GOLM1, because its transcript levels remain unchanged, this finding implies a post-transcriptional effect of cytokines to stabilize GOLM1. Following PERK or ISR inhibition, the relative occupancy of *CD274* in polyribosomes showed no change despite the further increase in its encoded protein levels (**Supplemental Figure 4F**). Altogether, these findings are consistent with the stabilization of the PD-L1 protein by GOLM1.

To correlate our findings to T1D, we next interrogated GOLM1 levels in mouse and human tissues. No differences in GOLM1 levels were observed in 8-week-old female NOD mice compared to age- and sex-matched CD1 and NSG mice by immunofluorescence staining (**Figure 5A and B**). With advancing age, pre-diabetic female NOD mice exhibit a reduction in GOLM1 levels in β cells (**Figure 5C and D**), suggesting that its decline may impair an otherwise more robust PD-L1 response. Upon treatment of 6-week-old NOD mice with HC5770 for two weeks, an elevation in GOLM1 levels was observed in β cells by immunofluorescence (**Figure 5E and F**) and in islets by immunoblot (**Supplemental Figure 4G**), although quantification of the immunoblot did not reach statistical significance. In human tissues, analysis of scRNA-seq data in the Human Pancreas Analysis Program (HPAP) showed that *GOLM1* mRNA increases in both quantity and in the proportion of β cells in individuals with single (N=8 donors) and double (N=2 donors) autoantibody-positivity and with T1D (N=9 donors) compared to non-diabetic controls (N=15 donors) (**Figure 5G**), suggesting that “surviving” β cells have more *GOLM1* mRNA. This increase in GOLM1 transcript is consistent with increases in *CD274* (encoding PD-L1) (**Figure 5G**).

**Figure 5:**
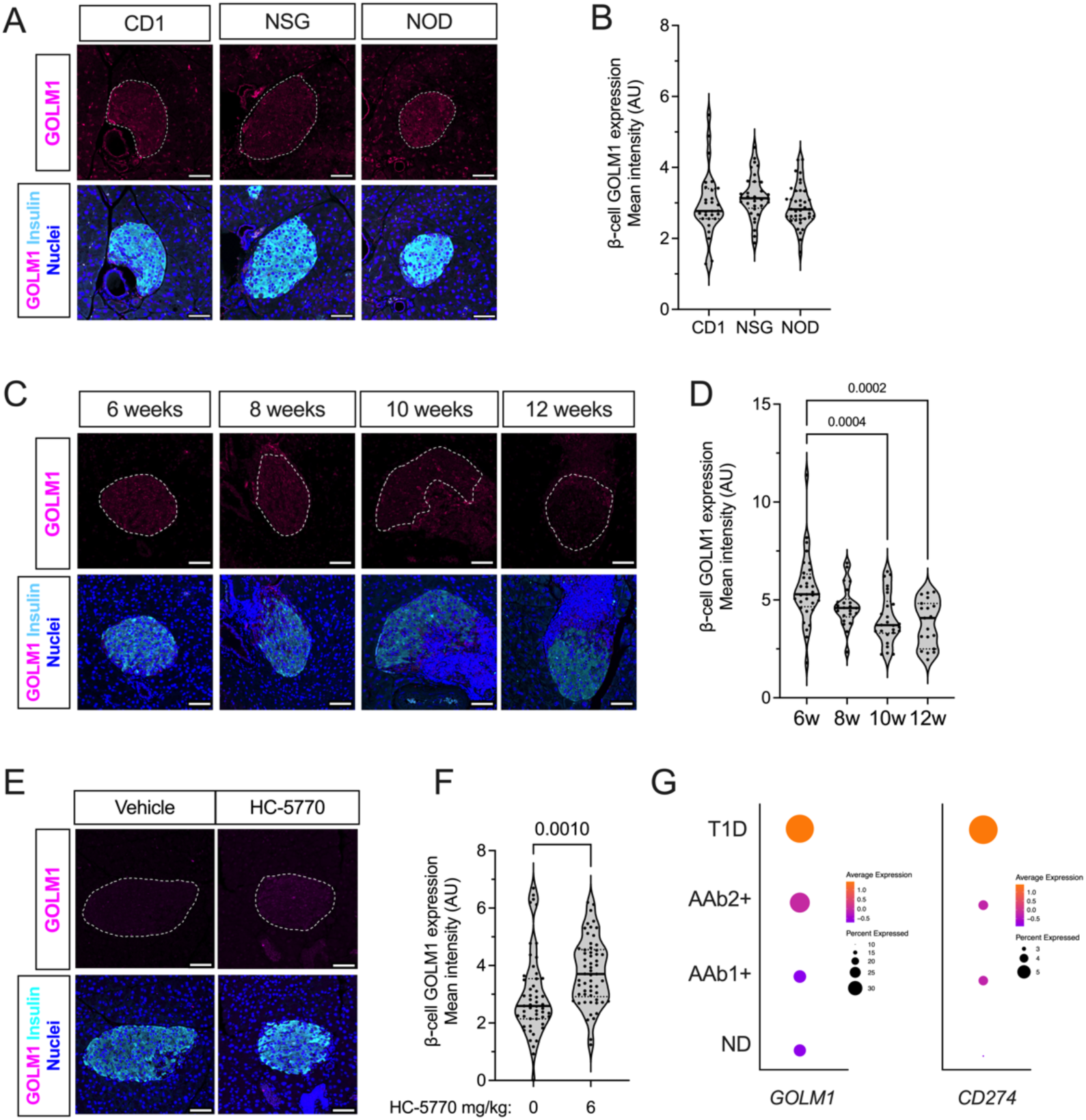
PERK inhibition increases GOLM1 levels in β cells. (***A***) Representative images of pancreata from 8-week-old female CD1, NSG, and NOD mice immunostained for GOLM1 (*magenta*), insulin (*cyan*), and nuclei (*blue*), scale bar = 50 μm; *dotted lines* indicate islet*s.* (***B***) Quantification of the GOLM1 fluorescence intensity from data in *panel (A)*; each dot represents an islet, N=3 mice, with >5 islets per mouse (ANOVA). (***C***) Representative images from 6-, 8-, 10-, and 12-week-old female NOD mice immunostained for GOLM1 (*magenta*), insulin (*cyan*), and nuclei (*blue*); scale bar = 50 μm; *dotted lines* indicate islet*s.* (***D***) Quantification of the GOLM1 fluorescence intensity from data in *panel (C)*; each dot represents an islet, N=3 mice, with >5 islets per mouse (ANOVA). (***E***) Representative images of pancreata from 8-week-old female NOD mice that have been treated with Vehicle of HC-5770 (6 mg/kg) for two weeks, immunostained for GOLM1 (*magenta*), insulin (*cyan*), and nuclei (*blue*), scale bar = 50 μm; *dotted lines* indicate islet*s.* (***F***) Quantification of the GOLM1 fluorescence intensity from data in *panel (E)*; each dot represents an islet, N=4-5 mice, and N>5 islets per mouse (T-test). (***G***) Dot plot analysis of scRNA-seq data in the Human Pancreas Analysis Program (HPAP) of residual β cells for *GOLM1* and *CD274*.The size of the dots indicates the percentage of cells that express the studied gene. The color scale shows the change of normalized and centered average gene expression within the different groups. No diabetes (ND): N=15; Single autoantibody positive (AAb1+): N=8; Double autoantibody positive (AAb2+): N=2; type 1 diabetes (T1D): N=9. Data are presented as mean ±SEM.

## DISCUSSION

With recent studies showing that targeting the immune system can delay T1D onset (1), the use of complementary approaches that target β cells raises the possibility of augmenting therapeutic efficacy to achieve more robust disease prevention. To date, such approaches remain limited, although some successes have been observed by targeting ER and oxidative stress (42, 43). Prior studies in mice (19–21, 44), human pancreas tissue (45, 46), and humans (47) suggest that ER stress in islet β cells contributes to both cellular dysfunction (reduced insulin secretion) and the production of neoantigens that trigger autoimmunity. The IRE1α and ATF6 arms of the ER stress cascade (for a review, see ref. (9)) have been genetically and/or chemically investigated in these prior studies, yet the role of PERK—an ISR kinase—has remained largely unexplored. In this study, we interrogated the PERK/ISR arm of the UPR pathway in the context of autoimmune diabetes. Because the genetic knockout of PERK in mice is known to result in endocrine and exocrine dysfunction during pancreas formation and maturation, we used a recently described PERK inhibitor, HC-5770 (30). Our results show that (a) inhibition of PERK during a period of β cell ER stress in NOD mice reduces insulitis, preserves β cell mass, and delays the development of diabetes, (b) gene expression patterns in β cells following PERK inhibition are consistent with reductions in the UPR and PERK response, and (c) inhibition of PERK activity augments the immune checkpoint protein PD-L1 through stabilization mediated by GOLM1.

In early T1D, β cells are exposed to inflammation, putative viral infections, hypoxia/ischemia, and/or impaired nutrient handling (resulting from insulin deficiency) (48–50). Moreover, genetic risk may increase the susceptibility of β cells to these underlying stress-inducing signals. For example, prior studies (and confirmed here) demonstrated that even under normal housing conditions, islets from immunodeficient NOD-SCID mice exhibit the UPR (44)— a finding suggesting that β cells of NOD strain may be especially prone to propagating inflammatory insults. Additionally, in humans, it has been observed that β cell death/stress and reduced pancreas mass are present in auto-antibody-negative, first-degree relatives of individuals with T1D (51, 52). In genetically high-risk infants, relative increases in glycemia are observed before the onset of seroconversion (53). These observations in humans support the possibility of a stressed state of β cells even in the absence of detectable autoimmunity. Under conditions of stress, β cells engage the ISR, an emergency response that is triggered by the activation of one or more of 4 kinases (PERK, HRI, PKR, GCN2), which act to reduce general protein synthesis and divert energy expenditure toward cellular recovery (54). Although it is adaptive in the short term, long-term ISR activation (or a greater magnitude of activation) potentially reduces the production of proteins necessary for cellular survival, thereby becoming maladaptive. A recent study demonstrated that the genes encoding 3 of the 4 ISR kinases (PERK, GCN2, PKR) are elevated in islets of T1D donors, with the gene encoding PERK (*EIF2AK3*) being notably dysregulated in islets of T1D donors (18). Similarly, in pancreatic tissue sections of autoantibody-positive donors, *EIF2AK3* is reported to be elevated compared to non-diabetic donors (18). These findings are also consistent with our own results here of elevated phosphorylated eIF2α (a proxy for ISR activation) and reduced protein translation in pre-diabetic NOD (and NSG) mice, collectively suggesting that the ISR might contribute to T1D development.

PERK activity in the pancreas is essential to support a functional β cell population, yet dysregulation and prolonged hyperactivation of PERK have been linked to several disorders, including cancer, diabetes, and neurodegeneration (55–57). This presents a challenge to interrogating PERK-driven disease: complete loss of PERK through genetic ablation or high-dose treatment with PERK inhibitors results in endocrine and exocrine pancreatic toxicity.

Previous studies have shown that genetic manipulations of PERK or eIF2α result in postnatal lethality and severe β cell deficiency (16, 58, 59). These findings may be related to activation of type 1 interferon signaling in the developing pancreas (36) and the requirement for PERK in neonatal and postnatal β cell expansion (23). Considering these prior observations, genetic models pose limitations on testing a direct role of PERK in the context of disease, in which the timing, duration, and extent of PERK activity may be critical to pathogenesis. To sidestep this issue, we made use of a selective PERK inhibitor that has a highly stable PK profile in mice.

PK/PD analyses confirmed that lower doses of HC-5770 attenuate PERK without completely abolishing PERK activity. By working within this dose range (0.6-6 mg/kg, QD), we demonstrated the therapeutic benefit of PERK inhibition by delaying T1D onset in the NOD mouse model without observable negative impact on the pancreatic islet. These studies suggest a reasonable safety window can be achieved through a dosing regimen, highlighting new therapeutic potential for PERK-driven diseases.

A finding that was only evident upon spatial proteomics analysis was the enhancement of β cell PD-L1 protein levels following HC-5770 treatment. The effect of HC-5770 to increase PD-L1 levels is likely related to PERK-mediated phosphorylation of eIF2α since we observed that blockade of the p-eIF2α/eIF2B interaction with ISRIB has a similar effect. The role of β cell PD-L1 in dampening the autoimmune attack through its interaction with the receptor PD-1 on immune cells is a highly engaging topic in the context of T1D treatment and islet transplantation, with studies reporting that the PD-L1/PD-1 interaction suppresses the adaptive immune response (60–64). Conversely, in humans, the use of immune checkpoint inhibitors (which block this interaction) increases the incidence of T1D in genetically susceptible populations (65, 66), representing one of the more common immune-related adverse events associated with this therapy. Our studies show that blockade of PD-L1 using a monoclonal antibody accelerated diabetes onset in control NOD mice as expected. Notably, the levels of hyperglycemia were attenuated through the inhibition of PERK, emphasizing the role of PERK inhibition in enhancing β cell PD-L1 levels. Our current finding highlights that PERK/ISR can be manipulated to enhance β cell PD-L1 levels to attenuate autoimmunity. Although it was previously shown that the ISR might potentiate the post-transcriptional production of PD-L1 in cancer cells (67), the mechanism by which the ISR suppresses PD-L1 levels in other disease contexts remains largely unexplored. A recent study (39) suggested that GOLM1 may stabilize PD-L1 protein. Consistent with that study, we show here that PD-L1 levels are stabilized by GOLM1, likely through direct interaction and suppression of ubiquitination. Notably, this stabilization by GOLM1 is not a universal feature of proteins shuttled to the membrane, as we did not observe similar effects on HLA-I.

Previous studies have shown that PD-L1 levels are elevated in the residual β cells of donors with T1D (68) as a possible explanation for the persistence of these cells. Our analysis of the scRNA-seq dataset of the HPAP dataset (69) suggests that gene encoding GOLM1 (*GOLM1*) is similarly elevated in the residual β cells of donors with T1D. Collectively, our studies demonstrate an axis linking PERK/ISR to post-transcriptional suppression of GOLM1, which in turn stabilizes PD-L1 protein levels in β cells.

Some key limitations of our study should be acknowledged. First, because of the early neonatal lethality of *Eif2ak3-/-* mice and the challenges of generating timed, tissue-conditional deletions on the NOD background, our study utilized pharmacologic inhibition of PERK in mice in vivo. Although our scRNA-Seq studies are consistent with PERK inhibition in β cells, and the target specificity of HC-5770 across kinome has been previously established (30), they do not fully exclude the potential for off-target responses. Additionally, our findings do not rule out a role for PERK in other cell types (e.g., immune cells or exocrine cells) that contribute to T1D pathogenesis. These limitations also reveal a strength of our studies—namely, the systemic administration of a new pharmacologic agent (HC-5770), which provides context for how PERK inhibition might be leveraged in humans for the prevention/delay of T1D. A final limitation is that our studies do not directly address if and how other ISR kinases (PKR, GCN2, HRI) might contribute to T1D development and if additional inhibition of these other kinases might potentiate the responses we observed and thereby more completely prevent disease.

Collectively, our studies emphasize the need to consider a maladaptive role of PERK in β cells in the pathogenesis of T1D and how inhibition of PERK might provide an opportunity, either alone or in combination with immune-modulating agents, for disease modification in T1D.

## METHODS

### Sex as a biological variable

Our study mostly examined female mice, because type 1 diabetes in the NOD strain is more frequently observed in females. However, some data from male mice are included that parallel those seen in females, suggesting that the effects observed in females may be relevant to males.

### Animals and procedures

Mouse experiments were performed under specific pathogen-free conditions and maintained in 12 h:12 h light:dark cycle with free access to food and water as per protocols approved by the University of Chicago Institutional Animal Care and Use committee. CD1 mice were purchased from Charles River (Charles River #022), and NOD.*Cg-Prkdc^scid^ Il2rg^tm1Wjl^*/SzJ (NSG; Jackson Labs #5557) and NOD/ShiLtJ (NOD; Jackson Labs #1976) mice were purchased from Jackson Laboratories (Bar Harbor, ME). Pharmacokinetic (PK) studies using BALB/c mice were performed by a contract to Pharmaron.

For diabetes incidence, 6-week-old female NOD mice were orally gavaged with vehicle (0.5% methylcellulose) or 0.6, 2, or 6 mg/kg HC-5770 (30) for either 2 or 4 weeks. 6-week-old female NOD mice were injected intraperitoneally with vehicle (5% DMSO, 2% Tween 80, 20% PEG400, and saline) or 0.25 or 2.5 mg/kg ISRIB (*trans*-isomer) (MedChemExpress: HY-12495) for two weeks (70). Blood glucose was measured by the tail vein using a glucometer (AlphaTrak). For diabetes incidence, diabetes was classified as two consecutive blood glucose values greater than 250 mg/dL. At the end of each study, mice were euthanized, and tissue and blood were collected. To isolate islets, collagenase was injected into the pancreatic bile duct to inflate the pancreas before removal, as previously described (71). Briefly, a Histopaque-HBSS gradient was applied to the dissociated pancreas, followed by centrifugation at 900 *x g* for 18 min. The mouse islets were then removed from the center of the gradient and cultured in RPMI medium. Islets were handpicked and allowed to recover overnight before experimentation.

For experiments involving anti-PD-L1, 6-week-old female NOD mice were orally gavaged with 6 mg/kg HC-5770 for a total of 4 weeks. When the mice were 8 weeks old, a single intraperitoneal injection of neutralizing rat monoclonal antibody against PD-L1(dosage: 200 mg/kg; BioXCell; BE0101) or rat IgG2b isotype control (dosage: 200 mg/kg; BioXCell; BE0090) were administered, while continuing the HC-5770 treatment for an additional 2 weeks. Control 8-week-old female NOD mice were administered a single dose of 200 mg/kg of anti-PD- L1 intraperitoneally. Glucoses were monitored over the subsequent 16 days post-injection of anti-PD-L1 or IgG control.

PK analysis of HC-5770 in mouse plasma followed a methodology described elsewhere (30). HC-5770 was suspended in a vehicle consisting of 0.5% methylcellulose (400 cP) and 0.1% Tween80 in water and administered to female BALB/c nude mice by oral gavage at 0.3, 1, 3, 10, 30 mg/kg. Plasma was sampled from 5 mice per group following a single oral administration at 1, 4, 8, 12, 24 h post-dosing. The plasma concentration of the compound was determined by protein precipitation with acetonitrile and liquid chromatography with tandem mass spectrometric detection (LC-MS/MS). Parameters were estimated using Phoenix (WinNonlin) pharmacokinetic software version 6.1.0 using a non-compartmental approach consistent with the oral route of administration. PK analysis in female NOD mice followed a similar methodology, with the exception that only the 1 and 10 mg/kg doses were evaluated. Methods describing the determination of mouse protein plasma binding were described previously (72). Frozen pancreata from mice treated with HC-5770 were homogenized and protein isolated in preparation for SimpleWestern as described previously (30). In brief: Protein detection was performed on the Jess SimpleWestern high-throughput protein analysis platform (ProteinSimple) according to manufacturer’s protocol using a 12-230 kDa Separation Module (ProteinSimple, SM-W004) and Total Protein Detection Module (ProteinSimple, DM-TP01). The following antibodies were used: p-PERK (Eli Lilly; 1:50) (73) and PERK (Cell Signaling Technology, 1:200, Cat. #3192).

### Human islets

De-identified non-diabetic male and female human donor islets were obtained from the Integrated Islet Distribution Program (IIDP) and the University of Alberta Diabetes Institute Islet core (**Supplemental Table 2**). The use of de-identified human samples was approved by the Institutional Review Board at the University of Chicago and considered exempt from human subjects research.

### Cell culture and treatment

MIN6 mouse β cells were cultured in high glucose (25mM) Dulbecco’s Modified Eagle’s Medium (DMEM) supplemented with 15% fetal bovine serum, 1% penicillin/streptomycin cocktail (P/S), and 1% L-Glutamine. Human EndoC-βH1 β cells (74) were cultured in low glucose DMEM (5.5mM) supplemented with 2% BSA, 50μM β-mercaptoethanol, 10 mM nicotinamide, 5.5 μg/ml transferrin, 6.7 ng/mL sodium selenite, and 1% P/S, in plates pre-coated with matrigel-fibronectin. HEK-293 cells were cultured in high glucose DMEM supplemented with 10% FBS, 1% P/S, and 1% L-Glutamine. Human islets were cultured in standard islet medium (Prodo) supplemented with human AB serum (Prodo), Glutamine and glutathione (Prodo), and ciprofloxacin (Fisher). Mouse islets were cultured in RPMI medium supplemented with 10% FBS and 1% P/S. Mouse Embryonic Fibroblast (MEF) cells were cultured in DMEM supplemented with 10% FBS.

Cells were pretreated with vehicle (DMSO), 250 nM HC-5770, or 50 nM ISRIB for 1 h followed by cotreatment with a proinflammatory cytokine cocktail for 18-24 h. For experiments involving MIN6 β cells, the proinflammatory cytokine cocktail contained 25 ng/mL mouse IL-1β (R&D Systems; 401-ML-010), 50 ng/mL mouse TNF-α (R&D Systems; 410-MT-010), and 100 ng/mL mouse IFN-γ (R&D Systems; 485-MI-100). For experiments involving human islets and EndoC-βH1 cells, the proinflammatory cytokine cocktail contained 1000 IU/mL human IFN-γ (R&D Systems; 285-IF-100) and 50 IU/mL human IL-1β (R&D Systems; 201-LB-005). EndoC- βH1 cells were transfected using Accell siRNA targeted against human *GOLM1* (Horizon Discovery). Experiments were performed 72-96 h post transfection for protein or 48 hours for RNA isolation. For experiments involving proteasome inhibition, 72 h post *GOLM1* knockdown, cells were concurrently treated with 10 μM MG132 and proinflammatory cytokine cocktail for 18- 24 h, after which protein was collected. For assessment of complementary activation of GCN2 upon PERK inhibition, islets were isolated from 9-week-old CD1 mice. After overnight recovery, islets were treated with increasing doses of HC-5770 (8nM-1μM), or 10μM harmine (Sigma; SMB00461-100MG) or DMSO for 24 h. At the end of the treatment, protein lysates were collected. In addition, protein lysates prepared from the whole pancreas of Balb/C mice were used as controls. MEF cells were treated with 1 μM thapsigargin (Sigma; T9033-1MG), 100 nM halofuginone (Cayman Chemicals; 13370), or DMSO for 6 h. At the end of the treatments, protein lysates were collected.

### Protein isolation and immunoblotting

Protein was isolated and western blots were performed as previously described (75). Briefly, whole-cell extracts of cells were prepared in a lysis and extraction buffer (ThermoFisher) supplemented with HALT protease inhibitor cocktail (ThermoFisher) and protein extract was resolved by electrophoresis on a precast 4-20% tris-glycine polyacrylamide gels (Bio-Rad), transferred to polyvinylidene difluoride membrane, and membranes were blocked with Intercept® (TBS) blocking buffer (Li-Cor Biosciences) for 1-2 h. The blots were probed with the following primary antibodies and 0.2% Tween20 with overnight incubation at 4°C: anti-p-eIF2α (Abcam; ab32157; 1:1000)(Cell Signaling; 3398s; 1:1000), anti-total-eIF2α (Cell Signaling; 2103; 1:1000), anti-puromycin (Millipore; MABE343; 1:5000), anti-ubiquitin (Cell Signaling; 43124; 1:1000), anti-HLA-I (ProteinTech; 15240-1-AP; 1:1000), anti-β-actin (Cell Signaling; 4970s; 1:1000)(Cell Signaling; 3700s; 1:1000), anti-PD-L1 (Cell Signaling; 29122s or 13654s; 1:1000)(Cell Signaling; 13684; 1:1000), anti-GOLM1 (Novus Biologicals; NBP1-50627; 1:1000), anti-pPERK (Cell Signaling; 3179; 1:500), and anti-PERK (Cell Signaling; 3192s; 1:500), anti-p- GCN2-T988 (Abcam; ab75836; 1:1000), anti-GCN2 (Cell Signaling; 3302s; 1:1000). Anti-rabbit or anti-mouse (Li-Cor BioSciences; 1:10000) or anti-Rabbit IgG (H+L)-HRP Conjugate (Bio-Rad; 170-6515; 1:5000) secondary antibodies were used for visualization and quantification. Immunoblots were visualized using the Li-Cor Odyssey system (Li-Cor Biosciences) and quantitated using Odyssey Imaging software (Li-Cor Biosciences) or ImageJ.

### Co-immunoprecipitation

Lipofectamine-based transfections of 25 μg pEGFP-PD-L1 (Addgene) and GOLM1 (Origene) vectors were performed in HEK-293 cells. To determine ubiquitination levels, HEK- 293 cells were transfected with plasmid pEGFP-PD-L1 24 h post-*GOLM1* knockdown. 24 h later, cells were treated ±MG132 overnight. 48 h post transfections, cells were washed, homogenized, and centrifuged. The clarified supernatant was incubated with protein A/G-agarose suspension (Santa Cruz) for 3 h at 4°C on a rocker to reduce background and remove non-specific adsorption of proteins. The supernatant was then incubated with either anti-PD-L1 (Cell Signaling 13684; 1:50) or IgG (Santa Cruz sc2027; 1:50). The supernatant was then resolved using SDS-PAGE gel as described above.

### Polyribosomal Profiling

Polyribosome profiling (PRP) experiments proceeded as previously described (76) with minor modifications. Briefly, 50 μg/mL cycloheximide was added to treated cells (in 100 mm plates) for 10 min to halt translation. Following cycloheximide treatment, cells were washed with ice-cold PBS containing cycloheximide and then collected in cell lysis buffer containing 50 μg/mL cycloheximide, 20 mM Tris-HCl (pH 7.5), 100 mM NaCl, 10 mM MgCl2, 1% Triton X-100, and 50 U/mL RNAse inhibitor and homogenized through 25 G needle. 10% input from the cytoplasmic supernatant was stored in RLT plus buffer with β-mercaptoethanol. The remaining supernatant was layered on a linear sucrose gradient of decreasing concentration (50% - 10%) and ultracentrifuged using a SW41Ti swing bucket rotor at 40,000 rpm for 2 h at 4°C. A piston gradient fractionator (BioComp Instruments) was used to fractionate the gradients, and absorbance of RNA at 254 nm was recorded using an in-line ultraviolet monitor. Total RNA from the PRP fractions was reverse-transcribed and subjected to SYBR Green I–based quantitative RT-PCR. P/M ratios were quantitated by calculating the area under the curve (AUC) corresponding to the polyribosome peaks (more than two ribosomes) divided by the AUC for the monoribosome (80S) peak.

### SUrface Sensing of Translation (SUnSET) assay

New protein production was determined using the SUnSET technique (28). Briefly, at the end of the treatments, cells were incubated with 10 μg/mL puromycin for 15 min for MIN6 cells and 30 min for mouse islets, and protein was isolated using RIPA lysis buffer and used for immunoblotting. After detection of puromycin, the blots were stripped using RESTORE™ plus stripping buffer (ThermoFisher; 46430) and then stained with REVERT™ 700 total protein stain (Licor; 926-11016). Puromycin lanes were normalized against corresponding lanes in total protein stain to account for loading differences.

### Immunofluorescence staining and quantification

Pancreata were fixed in 4% paraformaldehyde, paraffin-embedded, and sectioned. For immunofluorescence staining, pancreata were stained for using the following antibodies: anti-p- eIF2a (Cell Signaling; 9721; 1:200), anti-PD-L1 (Abcam; ab213480; 1:200), anti-PCNA (Santa Cruz; sc-7907; 1:100), anti-CD3 (Abcam; ab16669;1:100), anti-B220 (Biolegend; 03201; 1:100), anti-GOLM1 (Novus Biologics; NBP1-50627; 1:250 for mouse tissues and 1:50 for human tissues), anti-glucagon (Santa Cruz; sc514592; 1:50) and anti-insulin antibody (Dako IR002; 1:4). Highly cross-adsorbed Alexa Fluor secondary antibodies (ThermoFisher; 1:500) were used. Nuclei were identified through DAPI staining (ThermoFisher). All images were collected using a Nikon A1 confocal microscope.

Mean fluorescence intensity measurements for immunostainings in the β cell area were automated using CellProfiler v4.1 (77). Background subtraction was performed by removing lower quartile intensity pixels from each channel for each image. Fluorescence intensities were quantified in regions of interest defined by insulin-positive area. For tissues that had uneven illumination due to varying tissue depth, illumination correction was applied prior to intensity measurements following the CellProfiler tutorial for illumination correction across all cycles using Gaussian smoothing method.

### Immunohistochemistry and quantification

Pancreata were fixed in 4% paraformaldehyde, paraffin embedded, and sectioned to 5μm thickness. At least 3 sections, 100μm apart were used per mouse and immunostained with anti-insulin (ProteinTech; 15848-1-AP; 1:200) followed by recognition using Immpress reagent kit peroxidase conjugated anti-rabbit Ig (Vector Laboratories), DAB peroxidase substrate kit (Vector Laboratories) and counterstained with hematoxylin (Sigma). β cell death was determined using terminal deoxynucleotidyl transferase dUTP nick end labeling (TUNEL) using HRP-DAB chemistry (Abcam) performed as per manufacturer’s instructions on at least 2 sections, 100μm apart per mouse. Images were collected using a Keyence BZ-X810 fluorescence microscope system (Keyence) and the number of TUNEL positive cells was assessed manually per islet. β cell mass was calculated by calculating insulin+ area and whole pancreas area (62) using BZ-X800 Analyzer. The percentage of immune cell infiltration was scored as follows: 1 = no insulitis, 2 = infiltrate <50% circumference, 3 = infiltrate >50% circumference, 4 = infiltration within islet (44).

### Serum Insulin measurement

Serum insulin levels were measured using an ultrasensitive Insulin Enzyme-linked immunosorbent assay (Mercodia 10-1249-01) following the manufacturer’s guidelines.

### Single-cell RNA-sequencing

Islets were submitted to the University of Chicago Genomics Facility for library generation using 10X Chromium Single Cell 3’ v3.1 as previously described (78). Approximately 12,800 cells were loaded to achieve 8,000 captured cells per sample to be sequenced.

Sequencing was performed on Illumina Novaseq 6000. Raw sequencing files were processed through the Rosalind (https://rosalind.bio/) pipeline with a HyperScale architecture (Rosalind).

Quality scores were assessed using the FastQC tool (79). Cell Ranger was used to align reads to the Mus musculus genome build GRCm38, count unique molecular identifiers (UMIs), call cell barcodes, and perform default clustering. After initial processing, raw RNA matrices from each sample were then analyzed for quality control, “ambient” mRNA was removed using SoupX V1.6.1 (80), and clustering utilizing Seurat (81) v4.3.0 in R v4.2.2. Basic filtering parameters included cells with unique features of minimum 200 and maximum 7500. Cells expressing less than 25 percent mitochondrial related genes were included. Cell cycle effect was regressed out using previously established methods in Seurat. After filtering, replicates from each condition were merged using standard SCTransform Seurat protocols using 3000 integration features.

After clustering, cells were visualized using Uniform Manifold Approximation and Projection (82) and color customized using ggplot2 (83). Marker genes were determined using ‘FindAllMarkers’ function (Wilcoxon rank-sum test) in Seurat. Contaminating endothelial and neuronal cells were distinct from the other clusters and were subsequently removed from the final analysis.

Gene Set Enrichment Analysis (GSEA) was performed using the msigdbr package (https://cran.r-project.org/web/packages/msigdbr/vignettes/msigdbr-intro.html) to determine lists of genes from the following gene sets: Hallmark Gene Set (Unfolded Protein Response, Inflammatory Response), GO Biological Processes (PERK mediated Unfolded Protein Response), and Reactome (Antigen processing- Ubiquitin-Proteasome degradation). Pseudo- bulk differential gene expression analysis was performed where gene counts in all the cells for each biological replicate was aggregated and pathways identified by Gene Ontology (84).

Previously published single cell transcriptional data (25, 26) deposited in GEO (GSE141786, GSE117770) were reanalyzed using the above-mentioned protocol using R. GSEA for the Hallmark unfolded protein response pathway was performed on the endocrine sub-population from 4-week, 8-week, and 15-week-old NOD islets from GSE141786 dataset, and from the β cell population from 8-week, 14-week, and 16-week-old NOD islets from GSE117770 dataset.

FASTQ files of 10X Genomics scRNA-seq data for human islets were downloaded from the data portal of the Human Pancreas Analysis Program (HPAP) (69) (https://hpap.pmacs.upenn.edu). The datasets were generated from pancreatic islets samples of the donors indicated in **Supplemental Table 3**.

Raw reads were processed with Cell Ranger V6.1.2 (85) for quality control, alignment, and gene expression quantification. The reads were aligned to the human genome reference (GRCh38). The “ambient” mRNA was removed using SoupX V1.6.1 (80), employing genes (*INS*, *GCG*, *SST*, *TTR*, *IAPP*, *PYY*, *KRT9* and *TPH1*) identified from the initial clustering by Seurat (81) as markers that represent the major cell types of human islets. The potential doublet cells were assessed and removed by scDblFinder V3.16 (86). The remaining cells were further filtered by the following criteria: number of genes detected >200 and <9000, percent of mitochondrial reads <25% and number of counts <10000. Next, the SCTransform function (87) implemented in Seurat software was used to normalize the counts by removing the effects of library depth and regressing out the variation from the mitochondrial reads’ ratio. The top 3000 variable genes were selected to perform the principal component analysis (PCA). Finally, software Harmony V0.1.1 (88) was employed to integrate all the samples. This integration was performed based on the top 50 PCA components, considering the donor identity and reagent kit batches as the primary confounding factors. scSorter V0.0.2 (89) that uses known marker genes of cell types was utilized to annotate the cell types in human islets. After the above-described analyses, a total of 10,167 β cells from all the donors were identified, including 4,730 cells from non-diabetic donors, 2,372 cells from single autoantibody-positive donors, 2,480 cells from double autoantibody-positive donors, and 585 cells from type 1 diabetic donors.

### NanoString spatial proteomics

Paraffin embedded pancreata were used for nanostring spatial proteomics analysis. Tissues were stained with morphology markers: AF-647 conjugated insulin (Cell Signaling; 9008s; 1:400) and nuclei marker (SYTO13). Tissues were hybridized using a pre-validated mouse GeoMx Immune cell panel (NanoString; GMX-PROCONCT-MICP) comprising of the following markers: PD-1, CD11c, CD8a, PanCk, MHC II, CD19, CTLA4, SMA, CD11b, CD3e, Fibronectin, Ki-67, CD4, GZMB, F4/80, CD45, PD-L1; housekeeping genes: Histone H3, S6, GAPDH; and IgG antibodies: Rb IgG, Rat IgG2a, and Rat IgG2b for background subtraction. All the markers were conjugated to unique UV-photocleavable oligos for indexing. At least 5-6 islets with insulitis were chosen as regions of interest (ROI) per mouse based on the morphology markers (insulin and nuclei). The ROIs were segmented into insulitic region and insulin+ region for each islet. Oligos from the segmented ROIs were photocleaved, collected in a 96-well plate, and reads were counted using nCounter (Nanostring). Analysis was performed using nanostring software. Scaling was performed to normalize for any differences in tissue surface area and depth. After scaling, reads were normalized to housekeeping markers and background was subtracted using IgG markers. Normalized counts were visualized as heatmaps using GraphPad Prism.

### RT-PCR analysis

RNA stored in RLT plus buffer with β-mercaptoethanol was extracted using RNAeasy Mini kit (Qiagen), and cDNA synthesis was performed using High-Capacity cDNA Reverse Transcription Kit (Applied Biosystems) according to manufacturer’s instructions. SYBR-green-based quantitative PCR was performed using Bio-Rad CFX Opus. Relative gene expression was calculated using the comparative threshold cycle value (Ct), and normalized expression (to *ACTB* levels) is shown relative to vehicle control (ΔΔCT). For RNA from polyribosome profiling fractions, relative gene expression for monosome and polysome fractions was calculated with reference to the input (2^(input^ ^Ct-monosome^ ^or^ ^polysome^ ^Ct)^). Primers for *ACTB* (forward: 5’- GCACTCTTCCAGCCTTCCTT-3’; reverse: 5’-AATGCCAGGGTACATGGTGG-3’)*, GOLM1* (forward: 5’-GGATGTCCTCCAGTTTCAGAAG-3’; reverse: 5’- CTGTTCCTTCACCTCCTTCATC-3’)*, CD274* (forward: 5’-CCAGTCACCTCTGAACATGAA-3’; reverse: 5’-ATTGGTGGTGGTGGTCTTAC-3’) (Integrated DNA Technologies).

### Statistical Analyses

All data are represented as mean *±* SEM. For comparisons involving more than two conditions, one-way ANOVA or repeated measures ANOVA (with Tukey post-hoc test or Dunnett’s post-hoc test) was performed. For comparisons involving only two conditions, a two-tailed student’s unpaired t-test was performed. Mantel-Cox log-rank test was performed to determine the difference between groups in the NOD diabetes outcome experiments. GraphPad Prism v10 was used for all statistical analysis and visualization. Statistical significance was assumed at p-value < 0.05.

## ACKNOWLEDGEMENTS

We would like to thank Kara Orr (Indiana University School of Medicine) and Sarida Pratuangtham (University of Chicago) for their technical assistance. This work was supported in part by National Institutes of Health grants R01 DK060581 (to RGM), U01 DK127786 and U01 DK127786S1 (to RGM and BJWR), R01 DK133881 (to EKS and RGM), R01CA219815 (to SAO), F31 DK134070 (to CMA), T32 AI153020 (to JRE), an investigator-initiated award from HiberCell (to SAT and RGM), JDRF postdoctoral fellowship (3-PDF-2023-1326-A-N) and Diabetes Research Connection awards (both to CM), and a Quad Summer Scholarship award (to JEW). This study utilized Diabetes Center core resources supported by National Institutes of Health grant P30 DK020595 (to the University of Chicago) and P30 DK097512 (to Indiana University) and utilized services of the University of Chicago Histology and Genomics Cores.

## AUTHOR CONTRIBUTIONS

CM, VC, SAO, MES, RGM, and SAT conceptualized the research; CM, FH, JRE, JEW, JBN, TN, AC, KTF, SN, CMA, SS, EN, XY, DLE, BJWR, KAS, and SAT performed investigation; RGM and SAT provided project supervision; CM, SAT, and RGM wrote the original draft; all authors contributed to discussion, edited the manuscript, and approved the final version of the manuscript.

## DATA AVAILABILITY

RNA sequencing data have been uploaded to the public repository GEO with accession number GSE245004 with reviewer private access token <u>sxenuuksnlahjgb</u>. Any additional information required to reanalyze the data reported in this paper is available from the corresponding author upon request.

