## Supplemental Figures for "Inhibition of the Eukaryotic Initiation Factor-2-α Kinase PERK Decreases Risk of Autoimmune Diabetes in Mice"

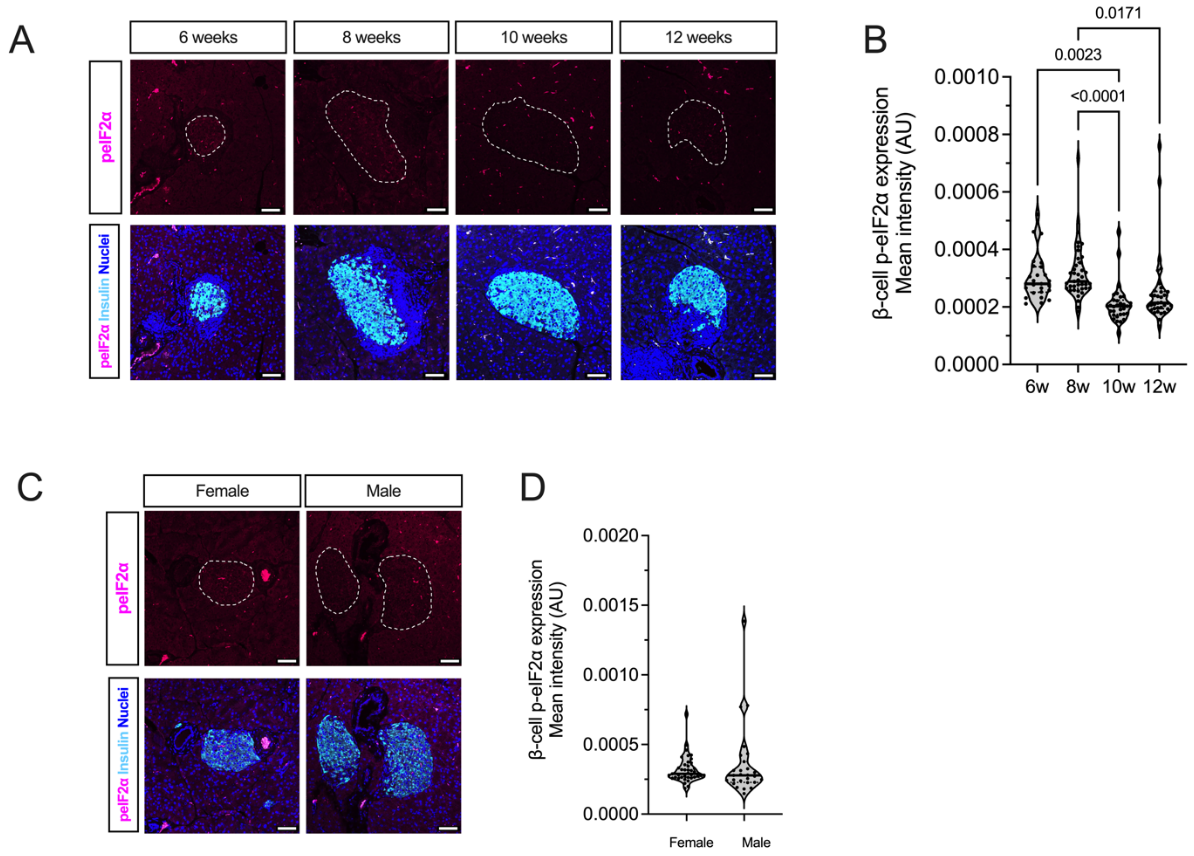


**Supplemental Figure 1: Phospho-eIF2α levels in NOD mice islets.** (***A***) Representative images of pancreata from female NOD mice at the indicated ages immunostained for phospho-eIF2α (p-eIF2α) (*magenta*), insulin (*cyan*), and nuclei (*blue*), scale bar = 50 µm; *dotted lines* indicate islets*.* (***B***) Quantification of the p-eIF2α fluorescence intensity from data in *panel (A)*; each dot represents an islet, N=4-5 mice, and N>5 islets per mouse (ANOVA). (***C***) Representative images of pancreata from 8-week-old male and female NOD mice immunostained for p-eIF2α (*magenta*), insulin (*cyan*), and nuclei (*blue*), scale bar = 50 µm; *dotted lines* indicate islets*.* (***D***) Quantification of the p-eIF2α fluorescence intensity from data in *panel (C)*; each dot represents an islet, N=4-5 mice, and N>5 islets per mouse (T-test).

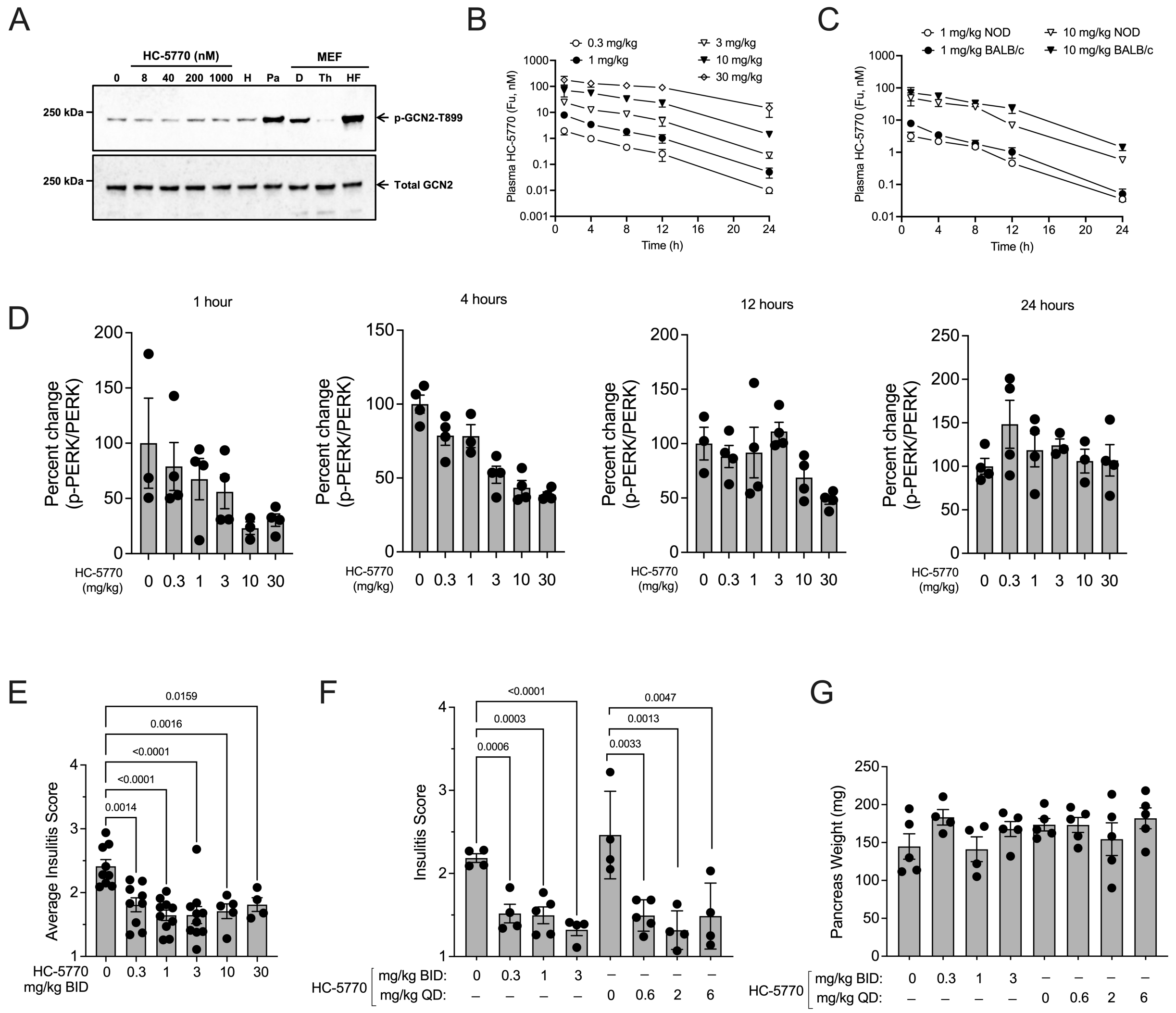


**Supplemental Figure 2: Pharmacokinetics and pharmacodynamics of HC-5770.** (***A***) Immunoblot analysis for phospho-GCN2 (p-GCN2) and total GCN2 of CD1 mouse islets, whole mouse pancreas (Pa), and mouse embryonic fibroblasts (MEF) under different treatment conditions: HC-5770, harmine (H), DMSO vehicle (D), thapsigargin (Th), or halofuginone (HF). (***B***) Free unbound concentration of HC-5770 in plasma following a single oral administration in BALB/c mice. HC-5770 quantified by LC-MS/MS (n=5 mice/group/time point). (***C***) Comparison of free drug concentrations in BALB/c and NOD mice plasma. HC-5770 quantified by LC-MS/MS following a single oral administration at 1 or 10 mg/kg. NB: BALB/c plasma data replicated from *panel (B)* for comparative purposes. (***D***) SimpleWestern® quantification of pPERK/PERK levels in BALB/c mouse pancreas following a single oral administration of HC-5770. (***E***) Insulitis scores in NOD mice following two weeks of twice daily (BID) treatment with HC-5770 (ANOVA). (***F***) Insulitis scores of NOD mice following two weeks of treatment with HC-5770 on a twice-daily (BID) or once-daily (QD) dosing schedule (performed contemporaneously). NB: QD dosing data are replicated in the data presented in Figure 2H, but they are shown here for comparative purposes (ANOVA). (***G***) Gross pancreas weight of NOD mice following two weeks of treatment with HC-5770 on both a QD and a BID dosing schedule (performed contemporaneously) (ANOVA). Data are presented as mean ±SEM.

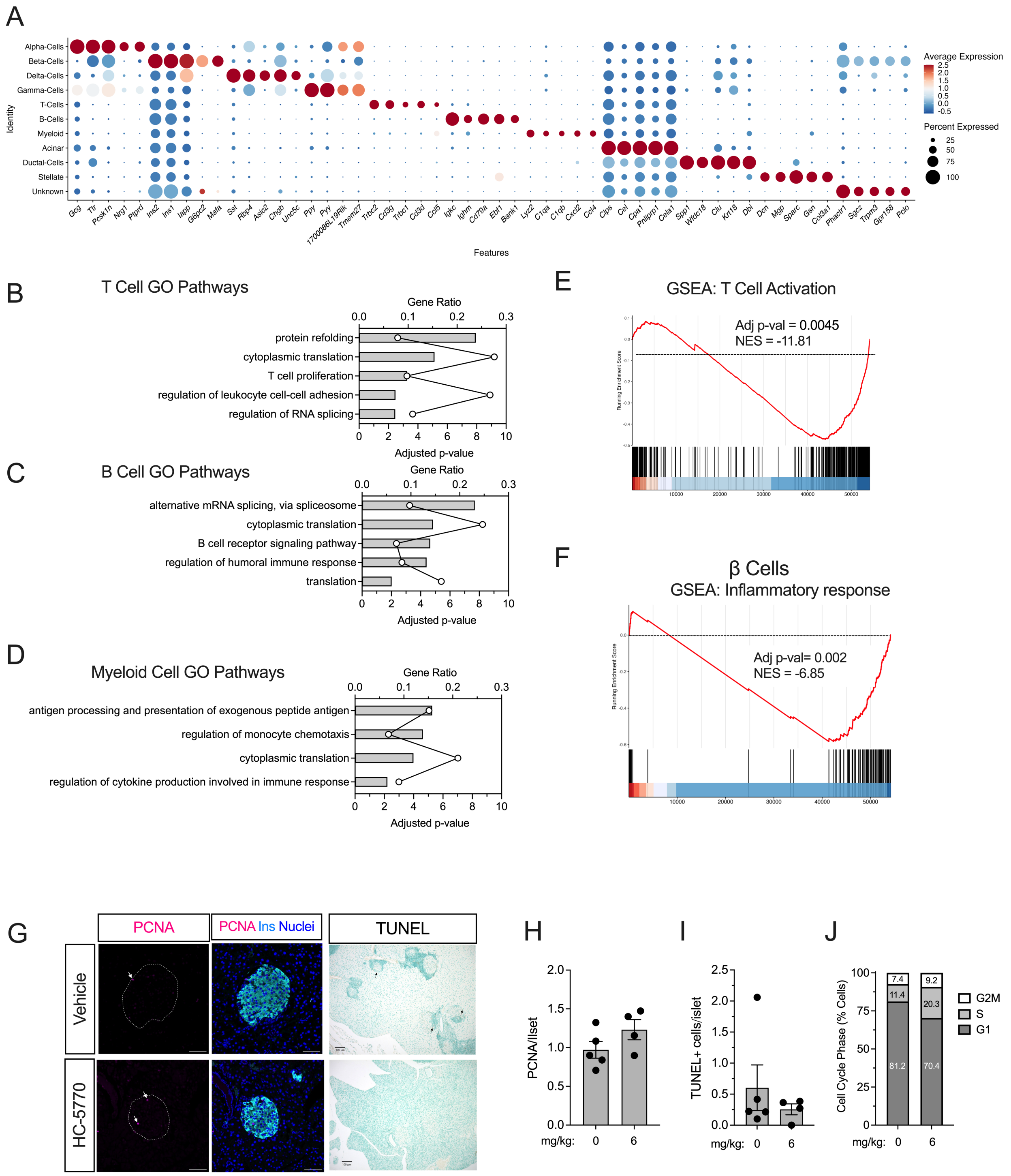


**Supplemental Figure 3. Single-cell RNA sequencing of islets from NOD mice treated with HC-5770.** 6-week-old female NOD mice were treated with Vehicle or HC-5770 (6 mg/kg) for 2 weeks, and isolated islets were subjected to scRNA-seq. (***A***) Dot plots of the top 5 genes used to identify cell type clusters. The size of the dots indicates the percentage of cells that express the indicated gene. The color scale shows the change in normalized average gene expression within the different groups. N=3 biological replicates for scRNA-seq. (***B)*** Gene ontology (GO) analysis of pathways observed in islet-resident T cells with HC-5770 treatment. (***C***) GO analysis of pathways observed in islet-resident B cells with HC-5770 treatment. (***D***) GO analysis of pathways observed in islet-resident myeloid-derived cells with HC-5770 treatment. (***E***) Gene set enrichment analysis (GSEA) of T cells showing HALLMARK: T cell activation. (***F***) GSEA of β cell clusters showing HALLMARK: Inflammatory response pathway. (***G***) Representative images of pancreata stained for PCNA (*magenta*, *arrows* indicate PCNA+ cells), insulin (*cyan*), and nuclei (*blue*); scale bar = 50 μm (*left and middle panels*) or TUNEL (*brown*, *arrows* indicate TUNEL+ cells), scale bar = 100 μm (*right panel*). (***H***) Quantification of PCNA+ cells in the islet; each data point represents the average of PCNA+ cells/islet from 3 separate pancreas sections from one mouse; N=4-5 mice (T-test). (***I***) Quantification of TUNEL+ cells in the islet; each data point represents the average of PCNA+ cells/islet from 3 separate pancreas sections from one mouse; N=4-5 mice (T-test). (***J***) Percentage of β cells in G1, S, and G2M cell cycle phases from scRNA-seq data. Data are presented as mean ±SEM.

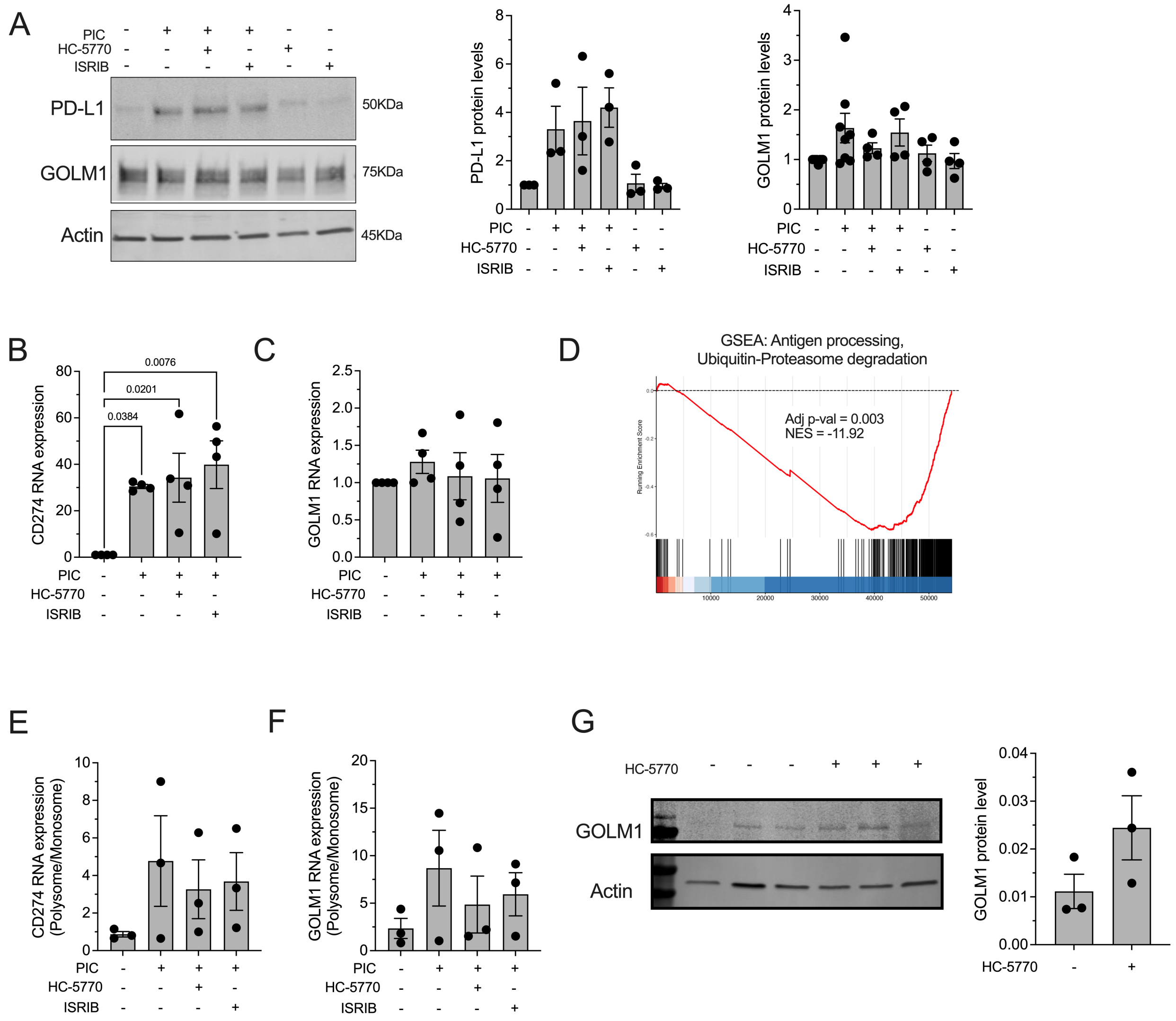


**Supplemental Figure 4. Coordinate increases in PD-L1 and GOLM1 in human islets and β cells and mouse islets.** (***A***) Representative immunoblot analysis of PD-L1 and GOLM1 of human islets treated with or without proinflammatory cytokines (PIC), HC-5770, or ISRIB (*left panel*); immunoblot quantification of PD-L1 levels (*middle panel*) and GOLM1 levels (*right panel*). N=3-8 independent experiments from different islet donors (ANOVA). (***B, C***) Relative *CD274* or *GOLM1* mRNA levels in human islets treated ±PIC, HC-5770, or ISRIB; N=4 independent experiments from different islet donors (ANOVA). (***D***) Gene set enrichment analysis (GSEA) of β cell clusters from scRNA-Seq following treatment of NOD mice with HC-5770 (Figure 3) showing REACTOME: antigen processing, ubiquitin-proteasome degradation. (***E, F***) Relative *CD274* and *GOLM1* levels in polyribosome to monoribosome fractions in human islets treated ±PIC, HC-5770, or ISRIB. N=3 independent experiments from different islet donors (ANOVA). (G) Immunoblot for GOLM1 and actin (loading control) from islets of 8-week-old female NOD mice treated with Vehicle or HC-5770 (6 mg/kg) for two weeks (*left panel*) and quantitation of the data from islets from N=3 mice per group (T-test). Data are presented as mean ±SEM.

**Supplemental Table 1: Pathway analysis for specific β cell clusters**

| Clusters | GO term | GO # | Gene ratio | -log10(adj p-val) |
|---|---|---|---|---|
| β0 | chaperone-mediated protein folding | GO:0061077 | 0.077 | 1.921 |
| | digestion | GO:0007586 | 0.056 | 1.521 |
| | cellular response to calcium ion | GO:0071277 | 0.055 | 1.526 |
| | translation | GO:0006412 | 0.029 | 2.500 |
| | regulation of apoptotic process | GO:0042981 | 0.013 | 1.467 |
| β1 | insulin secretion | GO:0030073 | 0.117 | 4.003 |
| | peptide hormone secretion | GO:0030072 | 0.094 | 4.068 |
| | antigen processing and presentation | GO:0019882 | 0.056 | 2.660 |
| | protein localization to extracellular region | GO:0071692 | 0.051 | 2.492 |
| | regulation of response to stress | GO:0080134 | 0.013 | 1.420 |
| β2 | cytoplasmic translation | GO:0002181 | 0.115 | 9.355 |
| | antigen processing and presentation of peptide antigen via MHC class I | GO:0002474 | 0.085 | 2.095 |
| | digestion | GO:0007586 | 0.067 | 2.357 |
| | response to endoplasmic reticulum stress | GO:0034976 | 0.035 | 2.032 |
| | regulation of apoptotic process | GO:0042981 | 0.013 | 2.040 |
| β3 | antigen processing and presentation of exogenous peptide antigen | GO:0002478 | 0.089 | 1.907 |
| | digestion | GO:0007586 | 0.078 | 2.592 |
| | secretion | GO:0046903 | 0.020 | 1.435 |
| | proteolysis | GO:0006508 | 0.014 | 1.839 |
| β4 | chaperone cofactor-dependent protein refolding | GO:0051085 | 0.152 | 3.264 |
| | digestion | GO:0007586 | 0.067 | 3.116 |
| | proteolysis | GO:0006508 | 0.014 | 3.827 |
| | response to stress | GO:0006950 | 0.007 | 1.678 |
| β5 | digestion | GO:0007586 | 0.089 | 4.678 |
| | response to hormone | GO:0009725 | 0.017 | 1.466 |
| | proteolysis | GO:0006508 | 0.013 | 1.613 |
| β6 | cytoplasmic translation | GO:0002181 | 0.044 | 2.212 |
| | proteolysis | GO:0006508 | 0.009 | 1.742 |
| β7 | response to endoplasmic reticulum stress | GO:0034976 | 0.026 | 1.759 |
| | protein processing | GO:0016485 | 0.024 | 1.313 |
| | secretion | GO:0046903 | 0.014 | 1.597 |
| | proteolysis | GO:0006508 | 0.014 | 5.539 |
| β8 | digestion | GO:0007586 | 0.044 | 1.757 |
| | proteolysis | GO:0006508 | 0.009 | 3.110 |

**Supplemental Table 2: Non-diabetic human donor islet characteristics**

| Donor ID | Islet source | Age (yrs) | Sex | BMI | HbA1c (%) |
|---|---|---|---|---|---|
| *RRID: SAMN19470079* | Integrated Islet Distribution Program (IIDP) | 39 | M | 27.6 | 5.5 |
| *RRID: SAMN19591106* | | 61 | M | 29.3 | 5.9 |
| *RRID: SAMN19897466* | | 28 | F | 24.7 | 5 |
| *RRID: SAMN28867622* | | 36 | M | 29.6 | 5.4 |
| *RRID: SAMN29657580* | | 40 | M | 23 | 5.2 |
| *RRID: SAMN30648329* | | 54 | M | 21.5 | 5.2 |
| *RRID: SAMN30986138* | | 67 | M | 34.4 | 5.3 |
| *RRID: SAMN32537360* | | 51 | M | 25 | 5.3 |
| *RRID: SAMN19796386* | University of Alberta | 40 | M | 31.7 | 5.4 |
| *RRID: SAMN19859645* | | 58 | M | 27.4 | 5.5 |
| *RRID: SAMN29094695* | | 61 | F | 36.1 | 5.8 |
| *RRID: SAMN33456515* | | 48 | M | 24.9 | 5.8 |
| *RRID: SAMN34997905* | | 40 | M | 30.4 | 5.4 |
| *RRID: SAMN35795155* | | 50 | M | 33.8 | 5.8 |
| *NDRI 2304-02252* | National Disease Research Interchange | 27 | M | 31.22 | N/A |

**Supplemental Table 3: Human Pancreas Analysis Program (HPAP) donor characteristics**

| Donor ID | Donor type | Age (yrs) | Sex | BMI | HbA1C |
|---|---|---|---|---|---|
| *RRID: SAMN19776470* | Non-diabetic | 5 | F | 16.3 | 6.8 |
| *RRID: SAMN19776475* | | 3 | F | 12 | 5.3 |
| *RRID: SAMN22562815* | | 4 | M | 20.63 | 4.9 |
| *RRID: SAMN19776478* | | 8 | M | 16.82 | N/A |
| *RRID: SAMN19776465* | | 13 | M | 18.6 | 5.2 |
| *RRID: SAMN19776467* | | 23 | F | 16 | 5.2 |
| *RRID: SAMN19776457* | | 24 | M | 20.8 | 4.9 |
| *RRID: SAMN19842611* | | 25 | M | 23.96 | 5.6 |
| *RRID: SAMN22562810* | | 28 | F | 24.7 | 5 |
| *RRID: SAMN19776458* | | 31 | F | 32.71 | 4.4 |
| *RRID: SAMN19776466* | | 35 | M | 26.91 | 5.2 |
| *RRID: SAMN19842585* | | 33 | M | 32.89 | 5.6 |
| *RRID: SAMN19776468* | | 35 | F | 21.9 | 5.3 |
| *RRID: SAMN19776471* | | M | 35 | 23.98 | 5.4 |
| *RRID: SAMN19776453* | | F | 39 | 34.7 | 4.7 |
| *RRID: SAMN19776455* | Donors with single islet-specific autoantibody (AAb1+) | M | 18 | 24.3 | 5.5 |
| *RRID: SAMN19776460* | | M | 23 | 28.6 | 5.3 |
| *RRID: SAMN19776469* | | M | 13 | 18.34 | 5.7 |
| *RRID: SAMN19776476* | | F | 27 | 26.2 | 5.2 |
| *RRID: SAMN19776480* | | M | 29 | 37.2 | 5.4 |
| *RRID: SAMN19776481* | | F | 21 | 28.99 | 5.1 |
| *RRID: SAMN19842601* | | M | 19 | 23.1 | 5.6 |
| *RRID: SAMN19842621* | | M | 21 | 25.59 | 5.6 |
| *RRID: SAMN19776474* | Donors with single islet-specific autoantibody (AAb2+) | M | 15 | 24.07 | 5.9 |
| *RRID: SAMN25600003* | | M | 15 | 23.59 | 5.3 |
| *RRID: SAMN19776451* | Donors with T1D | M | 14 | 13.2 | N/A |
| *RRID: SAMN19776452* | | F | 13 | 21.4 | N/A |
| *RRID: SAMN19776454* | | F | 17 | 21.35 | 8.9 |
| *RRID: SAMN19776463* | | F | 10 | 16.3 | 9 |
| *RRID: SAMN19776485* | | M | 24 | 27.9 | 10.4 |
| *RRID: SAMN19842593* | | M | 24 | 16.98 | 13 |
| *RRID: SAMN19842600* | | F | 12 | 15.42 | 9.8 |
| *RRID: SAMN19842613* | | F | 12 | 18.5 | 13.3 |
| *RRID: SAMN19842616* | | F | 15 | 19.3 | 10.4 |
